# Correlation analysis of target selectivity and side effects of FDA-approved kinase inhibitors

**DOI:** 10.1101/2021.03.18.435943

**Authors:** Omer Bayazeid, Taufiq Rahman

## Abstract

Kinase inhibitors (KIs) represent a popular class of therapeutic agents and chemical probes but most of them tend to be polypharmacological. Receptor and non-receptor Tyrosine KIs can target more than 100 kinases simultaneously compare to other KIs. We here analyze the molecular targets of 41 U.S. Food and Drug Administration (FDA)-approved KIs. We chose 18 drugs (Tyrosine KIs) and sought out to evaluate their selectivity profile and engagement with a number of targets in vivo at clinically relevant doses. We also wanted to see whether there prevails any correlation between the target engagement profile and the reported side effects for specific KIs chosen as test cases. To explore all clinical targets of the 18 KIs, we considered the free (unbound) maximum serum concentration (C_max_) of each KI and only chose targets for which the cognate affinities lie within the reported free C_max_ values, thereby allowing plausible interaction in clinical doses. We retrieved the side effects of those KIs that is reported in the FDA adverse event reporting system. We illustrate how correlation analysis of target−side effect can give a new insight into the off target of KIs and their effect on increasing the toxicity of KIs. These analyses could aid our understanding of the structural-activity relationship of KIs.

## 1. INTRODUCTION

The main mechanism used by eukaryotic cells to respond to both intracellular and extracellular singles is protein phosphorylation which is controlled by protein kinases ^1^. There are over 518 kinases in human kinome and it has been estimated that they interact with up to 30% of all human proteins ^2^. Protein kinases are mainly classified into two groups; serine/threonine kinases and tyrosine kinases. Kinases play important role in cell proliferation, migration, survival and apoptosis ^3^. Any dysregulation/structural mutations in kinases or their signaling pathways can result in various diseases that notably include cancer. Protein kinases share common structural aspects in having two lobes; a small N-lobe (five-stranded β-sheet and α-helix called the C-helix) and a large C- lobe (six α-helices). The two lobes are connected by a hinge region forming the active site (ATP-binding site) ^4^. Most of anticancer kinase inhibitors (KIs) target the ATP-binding site based on the idea if ATP is not able to bind to the protein kinase, phosphorylation of the substrates cannot occur. The catalytic domain of protein kinases appear to be highly conserved across the whole kinome and this makes it very difficult to achieve desired level of kinase selectivity for the KIs. ^5^. In contrast, protein kinases have developed complex regulatory mechanisms, and this led to discover another class of KIs (allosteric inhibitors). This class binds to sites outside the catalytic domain and are less conserved across the kinome which may offer a solution for the kinase to have a higher selectivity ^6^. Additionally, active kinases have very similar conformation, unlike inactive kinases. This also can aid the development of more selective KIs. The ATP-bind site is highly conserved in all the kinases. There are three conserved motifs that are essential for catalysis in the ATP-binding site; AXK motif (Ala-X-Lys, located in β3 strand), DFG motif (Asp-Phe-Gly, located in the N-terminal), Y/HRD motif (His-Arg-Asp, located in β7 strand). Another important motif is the APE motif (Ala-Pro-Glu, located in the N-terminal) ^7^.

## 2. PROTEIN KINASE DOMAIN

### 2.1. Phosphate-binding loop (P-loop)

The P-loop only interacts with the non-transferable phosphate atoms of ATP. It contains glycine-rich loop and the AXK motif ^8^. Glycine-rich loop is highly conserved motif with a sequence of **Gly**-X-**Gly**-X-X-**Gly** where X can be any amino. Its located in the N-lope and it extends over the ATP and acts as seatbelt to hold the ATP in the activation site during the process of phosphorylation ^9^. Next to the Glycine-rich loop there is the AXK motif. The most important amino acid in AXK motif is **Lys**. In the active state, **Lys** form a salt bridge with **Glu** in the C-helix and subsequently binds to the ATP molecules by forming hydrogen bonds with the oxygen atom of **α** and **β** phosphates (**Figure 1**) ^10^.

**Figure 1.**
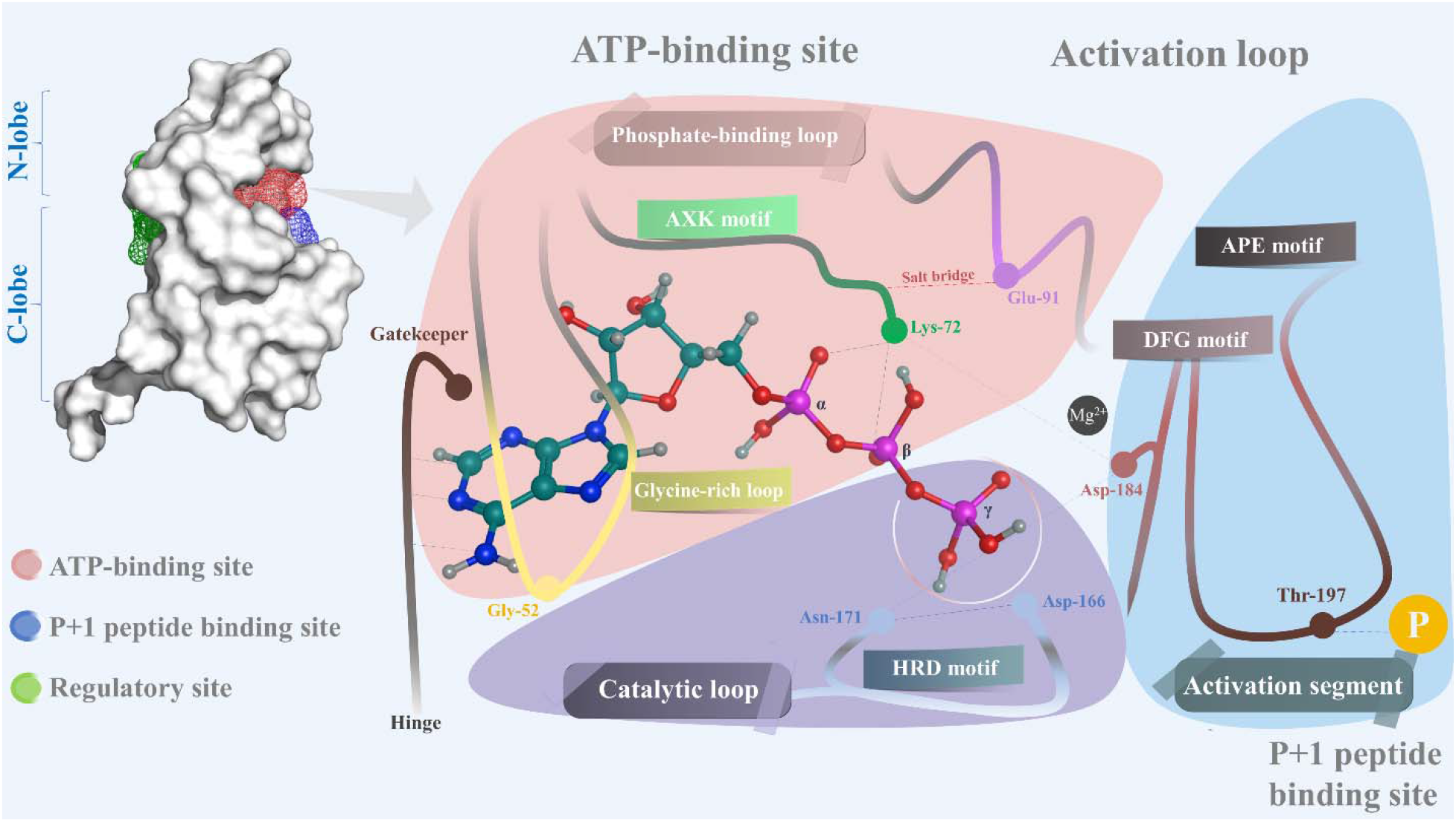
The ATP-binding site on kinases. The crystal structure (white), belongs to protein kinase A (3PLQ). Hydrogen bonds were retrieved from Knighton et al.^12, 13^.

### 2.2. Activation loop (A-loop)

The activation loop begins with the Asp-Phe-Gly (DFG) motif and ends with Ala-Pro-Glu (APE) motif (both highly conserved). The DFG motif has three residues, **Asp** (small in size), **Phe** (big), **Gly** (small). In the active form, the DFG motif sits inside the active site, this is known as “DFG-in” conformation. This allows the **Asp** of the DFG motif to bind to the magnesium ion (Mg^2+^) which is needed to transfer the γ phosphate. The magnesium ion catalyzes the phosphoryl transfer by binding to the oxygen atom of the **β** phosphate ^4^ (**Figure 1**). Some kinases require two Mg^2+^ ions. When the kinase is active, the APE motif recognizes the substrate and form a clef to bind to it. Close to the ATP-binding site, protein kinases have another binding site for the target protein to be phosphorylated. ^11^.

### 2.3. Catalytic loop

When the substrate binds to the activation loop, it interacts with the **Asp** of the Y/HRD motif in the catalytic loop. **Asp** catalyze the transfer of **γ** phosphate to tyrosine, serine or threonine of the substrate (**Figure 1**). Another important conserved part of protein kinase is glycine rich lope.

### 2.4. Gatekeeper

Behind the adenine ring binding site of ATP there is a back hydrophobic cavity that is not occupied by ATP. This gatekeeper blocks any molecules from occupying the cavity. Its size and shape depend on residue type which is the first residue of the hinge that connect the N-lobe with the C-lope. A smaller residue such as **Thr-529** in B-RAF allows the inhibitor to enter and bind to the back pocket (Figure 2A). When the residue is large (**Phe-80** in CDK2), it sterically blocks inhibitor from entering the back pocket (Figure 2B) ^14^. This is can be very effect to target specific kinases with small gatekeeper residue.

**Figure 2.**
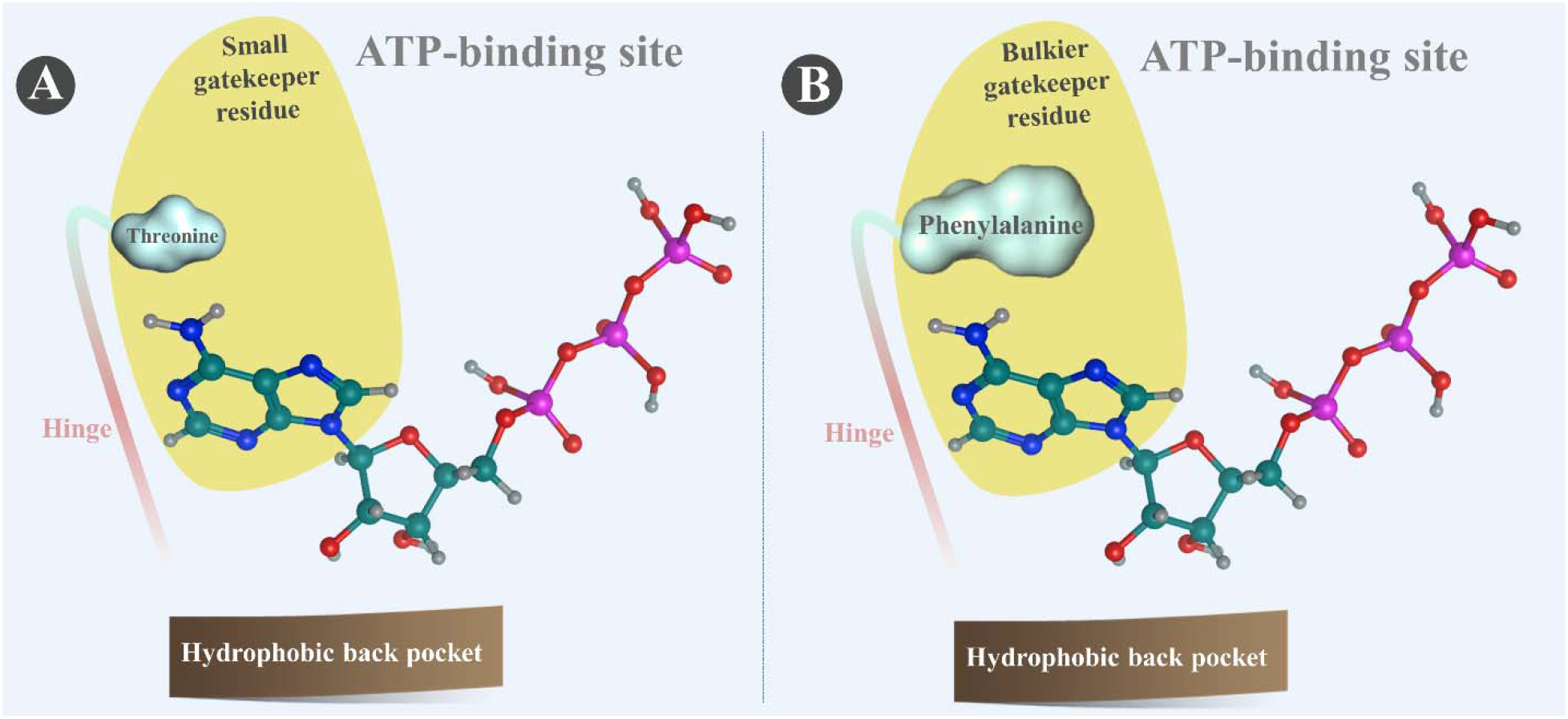
Gatekeeper in the hydrophobic back pocket of the ATP-binding site of Kinases.

### 2.5. Inactive kinase conformation

Kinases exhibit a completely different conformation when they are inactive. In inactive form, the activation loop spread out on the surface of the protein blocking the substrate from binding. In addition, the DFG conformation changes from “DFG-in” to “DFG-out” prevents the Mg^2+^ ion from interacting with ATP ^15^. In DFG-out state, the big residue **Phe** swaps place with the small **Asp** thus removing **Asp** from the activation site and block the entrance to the ATP-binding pocket ^16^.

## 3. TARGET ANALYSIS OF KIS

KIs have proven their success in treating several types of cancer. Protein kinases come second after the G-protein-coupled receptors as the most targeted group of proteins ^17^. A recent publication by Klaeger et al. showed that some KIs can target more than 100 protein kinases simultaneously while other KIs target fewer kinases^18^. We collected 41 U.S. Food and Drug Administration (FDA)-approved KIs from ChemBL database (https://www.ebi.ac.uk/chembl/) based on their single primary human targets ^19, 20^ and plot the targets with half-maximal inhibitory concentration (IC_50_), inhibition constant (K_i_) and mean dissociation constant (K_d_) with a cut off of <7 μM for the latter (Figure 3A). Additionally, we also collected the targets of the 41 KIs from DrugBank (https://go.drugbank.com/) ^21, 22^ and plot it in Figure 3B. There appears to be large differences between the target reported in ChemBL and the target reported in DrugBank. To explore the other clinical targets of these kinases, we decided to calculate the free (i.e. unbound to plasma proteins) maximum serum concentration (C_max_) of each KI and only chose targets for which the cognate affinities lie within the reported free C_max_ values, thereby allowing plausible interaction in clinical doses. According to the heatmap clustering, we decided to choose the KIs with the most targets (highlighted in Figure 3A) to enlist the targets these KIs are likely to engage with in vivo. The calculation is based on the highest C_max_ reported in the literature (**Table 1**). A heatmap showing KIs with few targets can be found in the supplementary file (**Figure S1**).

**Table 1.**
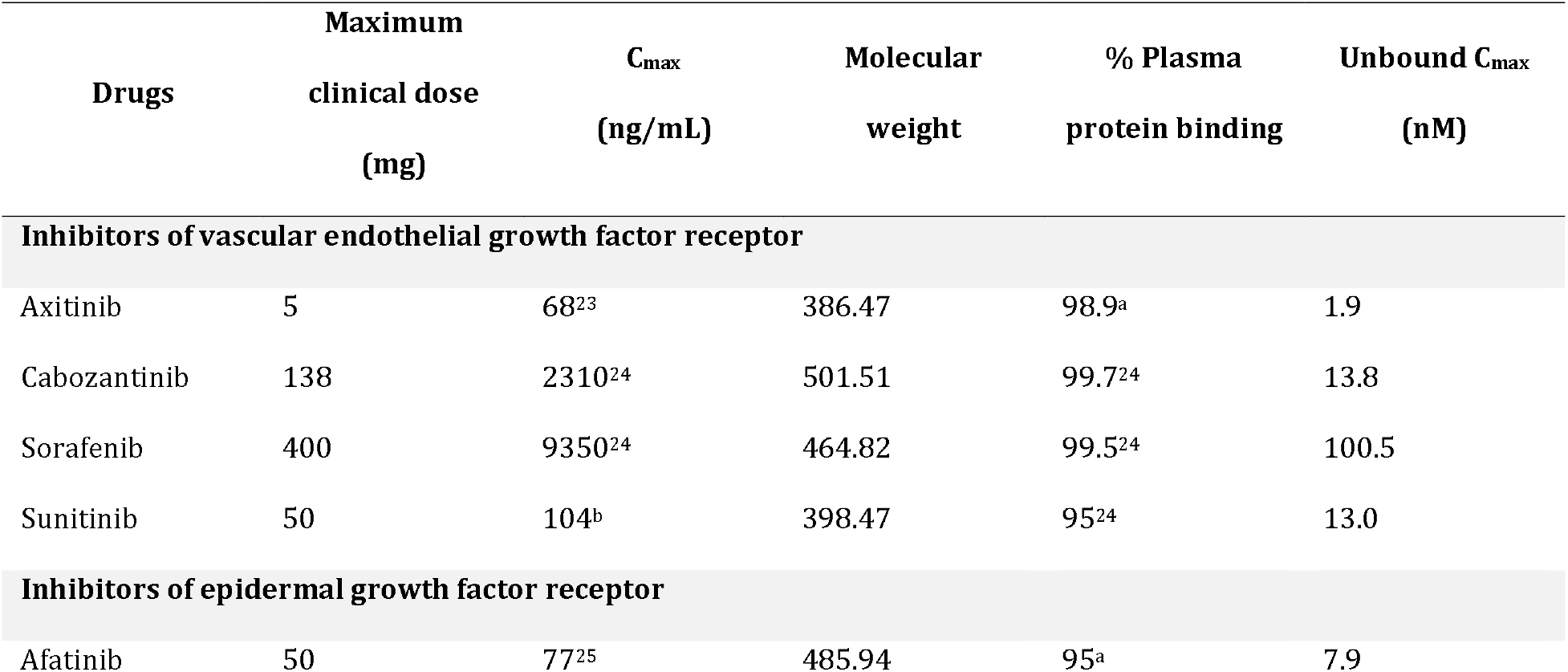
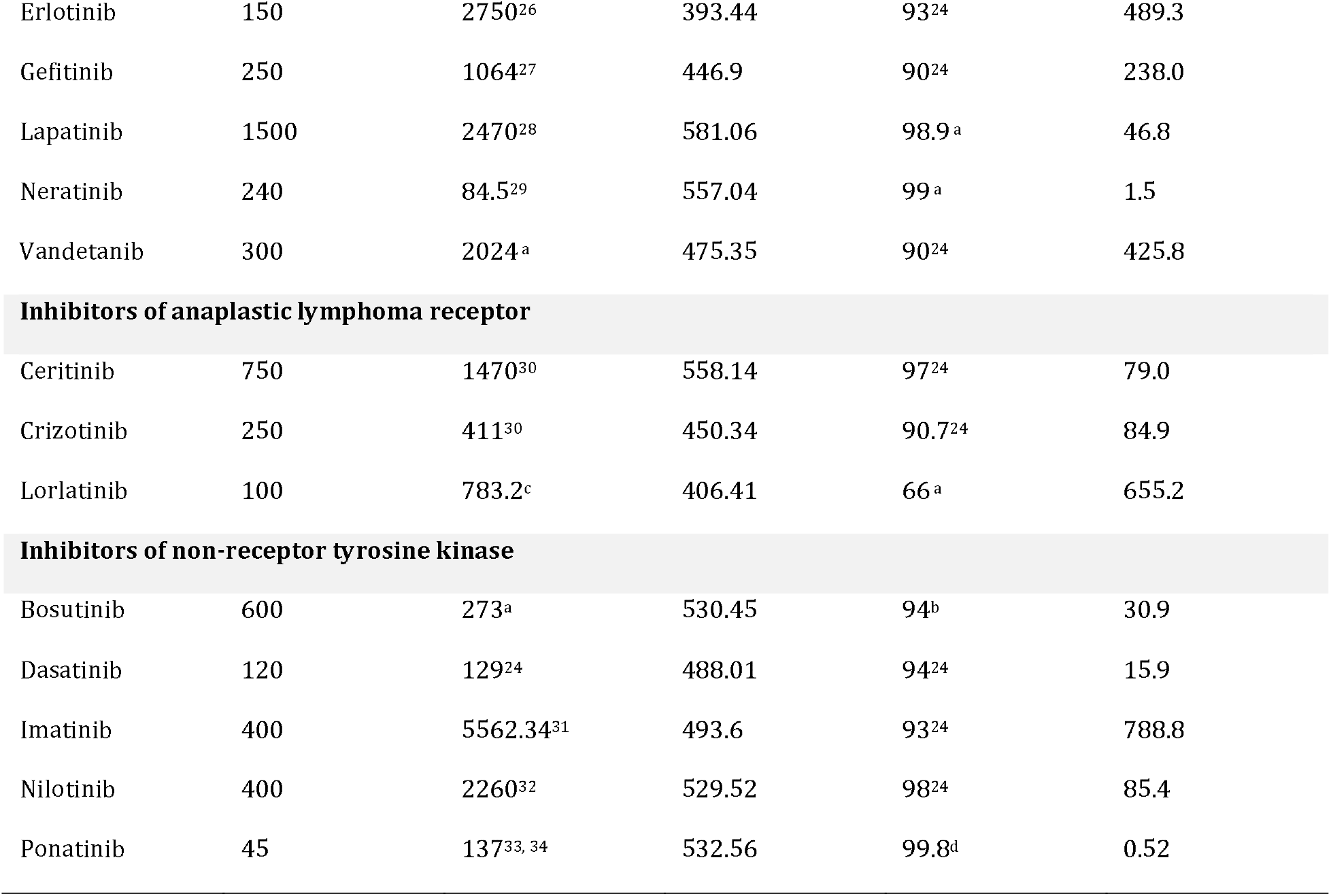
Unbound C_max_ of the KIs. ^a^FDA, ^b^European Medicine Agency, ^c^ Japanese Medicine Agency, ^d^Australian Medicine Agency.

**Figure 3.**
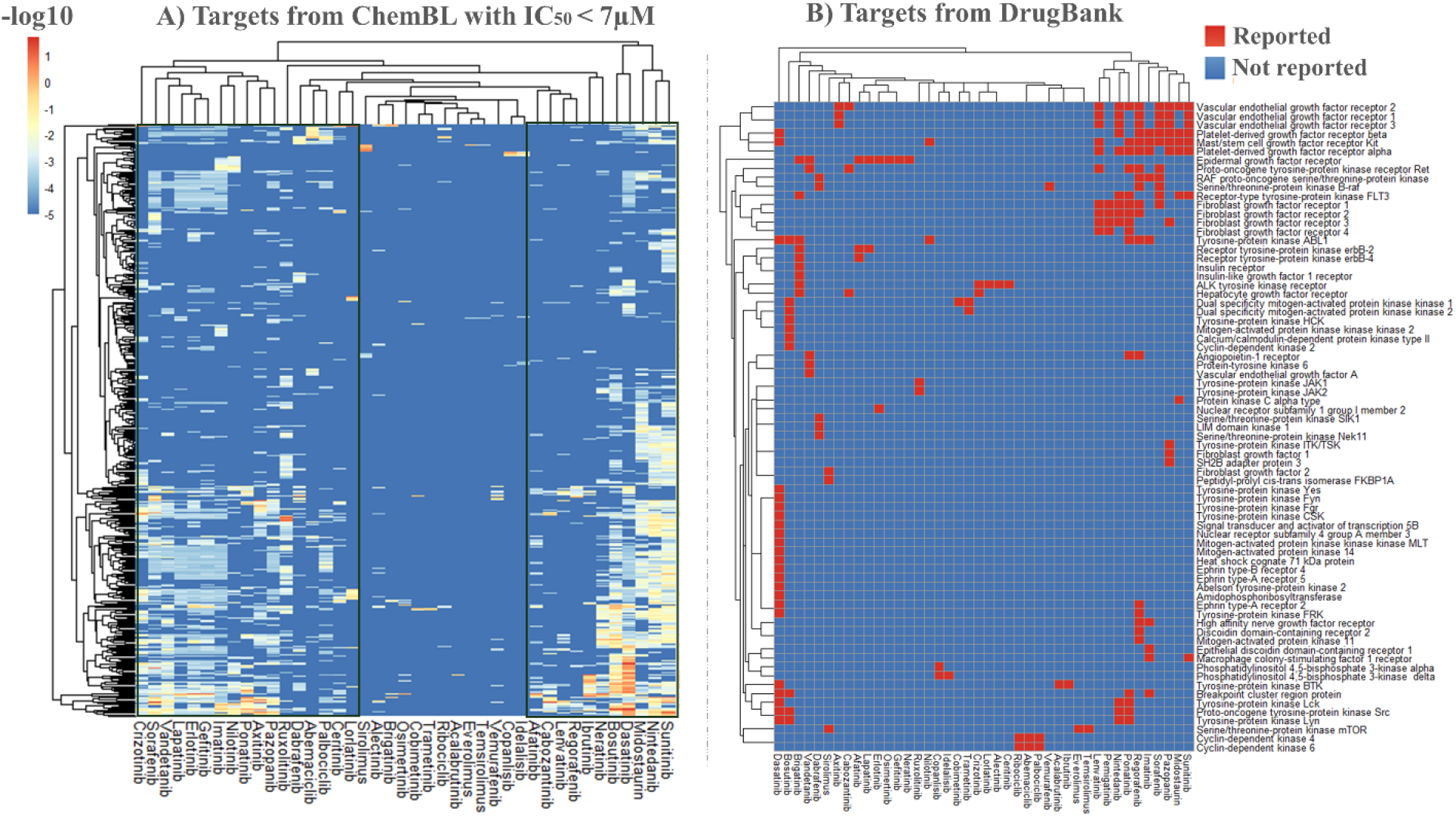
Heatmaps showing the targets of the 41 KIs. Values in (A) are showed in −log_10_(IC_50_,K_i_, K_d_).

### 3.1. Tyrosine kinase (TK)

TKs are enzymes that catalyze phosphorylation of a specific tyrosine residues in target proteins. TK signaling pathways often control cell proliferation and prevent any over proliferation of cells which could lead to the accumulation of abnormal cell numbers in the tissues/organ. Genetic or epigenetic alteration of TK signaling pathways result in forming and accumulating of cancer cells ^35^. Tyrosine kinase family is the most targeted family among the protein kinases, almost half of the KIs approved by the FDA, their primary target/s are tyrosine kinases. TKs are divided into two main classes; receptor TKs and non-receptor TKs. Receptor TKs facilitate communication between the cells and control cell growth, differentiation, motility, and metabolism. In total, there are 58 human TKs and they all share similar structure ^36^. Most growth factor receptors are TK receptors. There are 20 receptor TK classes, seven of these classes —epidermal growth factor receptor (EGFR), insulin receptor, platelet-derived growth factor receptors (PDGFR), fibroblast growth factor receptor (FGFR), Vascular Endothelial Growth Factor (VEGF), hepatocyte growth factor receptor (HGFR), and anaplastic lymphoma kinase (ALK) receptor — are highly associated with cancer ^37^. The targets of the selected 18 FDA-approved KIs have been retrieved from ChemBL database (https://www.ebi.ac.uk/chembl/) ^19, 20^. We filtered the targets and we only selected the targets within the range of free (i.e. unbound to plasma proteins) C_max_. We generated a heatmap for receptor TKs of the selected 18 KIs. All the heatmaps in this perspective are generated by using RStudio software (version 1.2.5042) ^38^. Out of the 18 KIs; 8 target the EGFR family, 1 targets insulin receptor, 6 target PDGFR family, 2 target FGFR family, 7 target VEGFR family and 2 target HGFR. Most of the KIs that target VEGFR also target PDGFR and stem cell growth factor receptor (**Figure 4**).

**Figure 4.**
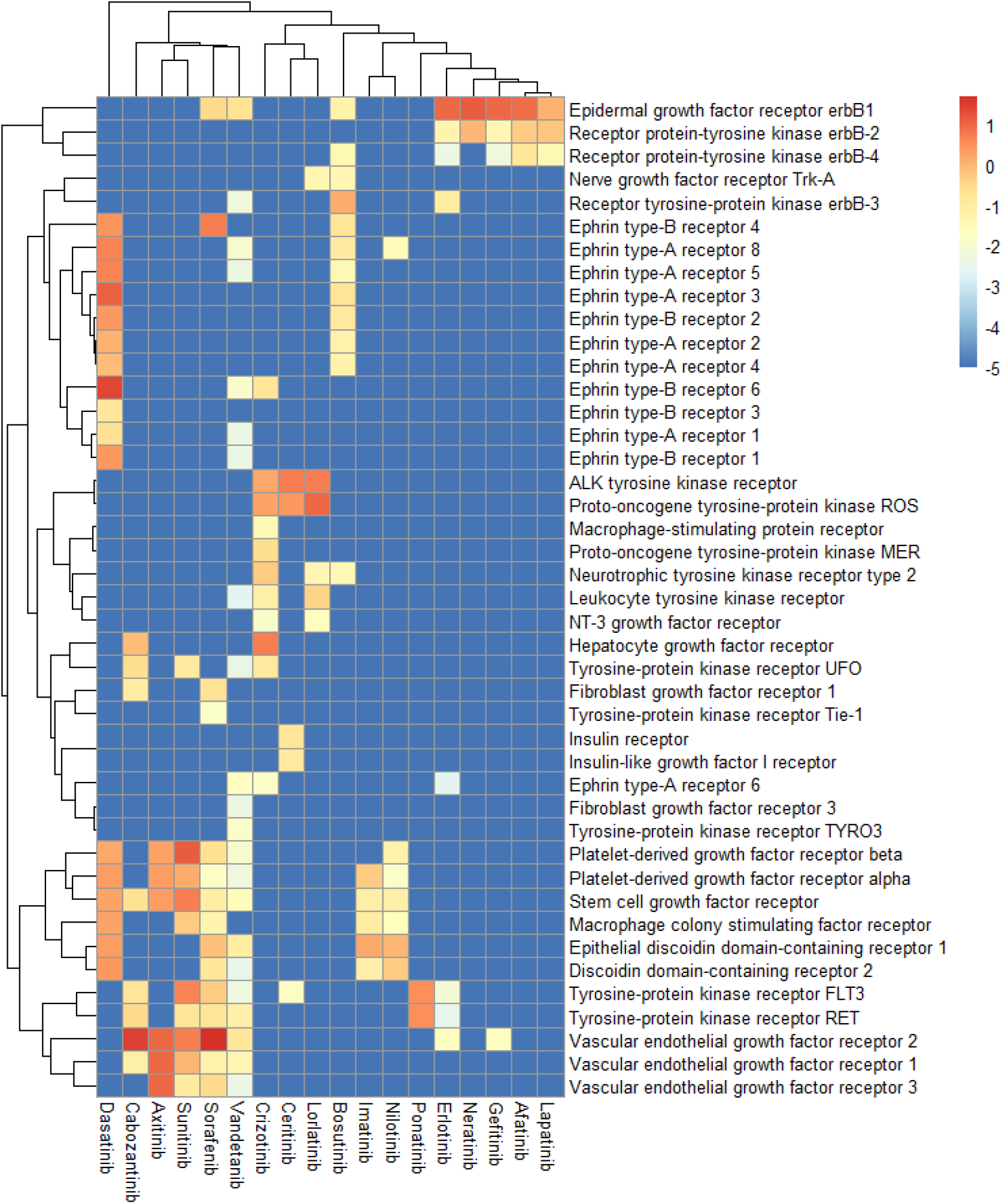
Tyrosine kinase receptors: the heatmap showing the targets of the 18 KIs (unbound C_max_). Values are showed in −log_10_(IC_50_, K_i_, K_d_).

Non-receptor TKs are enzymes that transmit signals inside the cells. They can either bind to the cell membrane or they can be nuclear-specific. Non-receptor TKs are divided into 9 families ^39^. The Abelson (ABL) kinase family regulates proliferation, migration, invasion and adhesion. Feline sarcoma (FES) kinases family takes part in cell cycle progression ^40^. Focal adhesion kinase (FAK) family regulates proliferation and cell adhesion. Activated Cdc42 kinases (Acks) family plays an important role in cell growth through stimulating Janus kinase (JAK) and SRC kinases ^41^. The targets of the selected 18 FDA-approved KIs have been retrieved from ChemBL and filtered according to the unbound C_max_. We generated a heatmap for the non-receptor TKs of the selected 18 KIs (**Figure 5**). In this perspective we will focus on three receptor TKs; VEGFR, EGFR and ALK receptor and the Non-receptor TK ABL.

**Figure 5.**
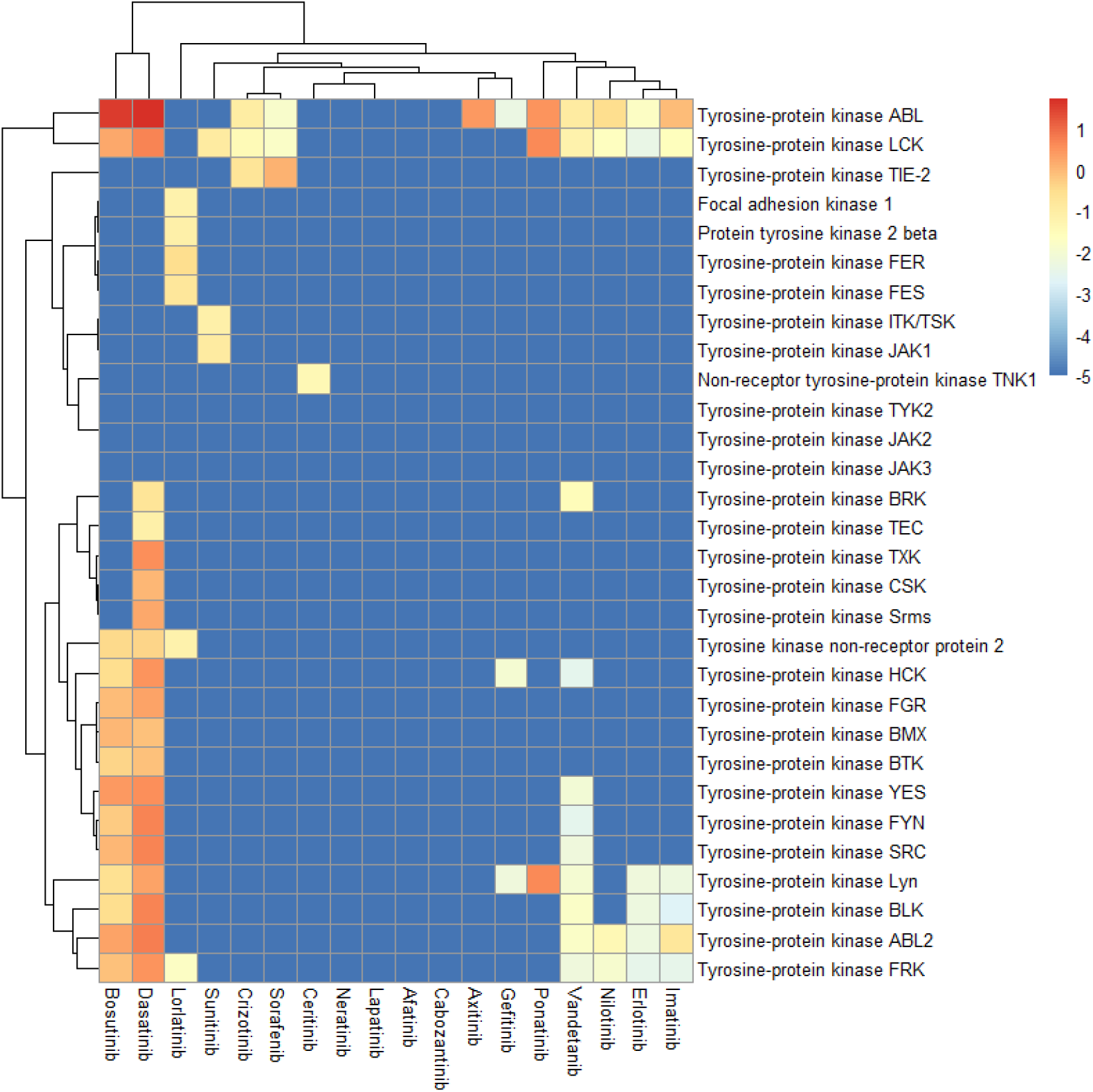
Non-receptor tyrosine kinase: the heatmap showing the targets of the 17 KIs (unbound C_max_). Values are showed in −log_10_(IC_50_, K_i_, K_d_)

To see if there is a correlation between the clinical targets of the selected KIs, we collected the side effect data from Target validation website (www.targetvalidation.org) ^42^. The side effects are retrieved from the FDA adverse event reporting system. The report values are in Log likehood (LL) ratio. We re-scaled the values to be between 1 and zero where 1 is the highest LL ratio and zero indicative of no reported side effect. We did this for each drug separately. Correlation analysis using Pearson’s test was computed using RStudio software ^38^(**Figure 6**). The full list of the KIs’ side effects can be found in the supplementary document (Table S1).

**Figure 6.**
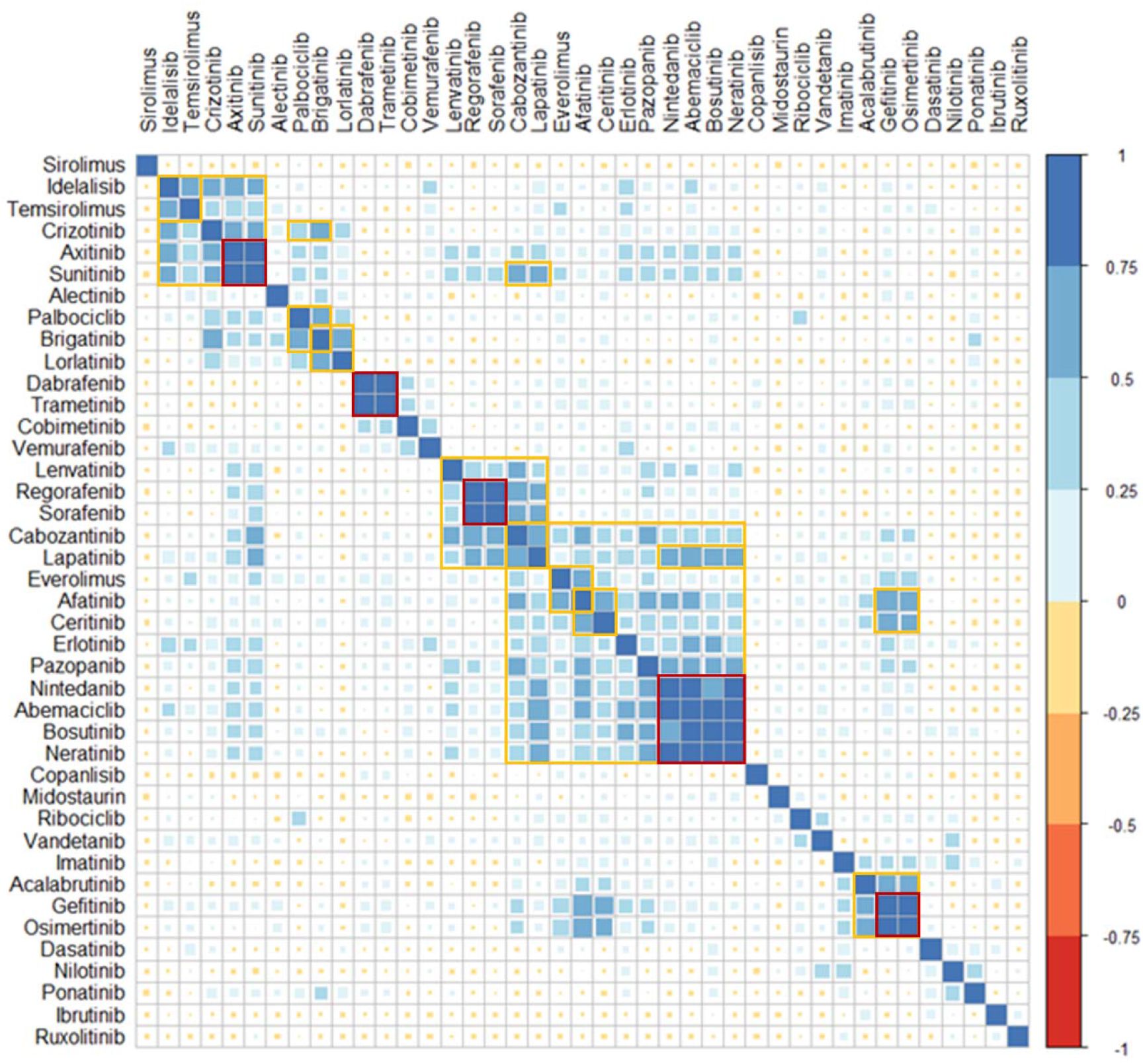
Correlation analysis of the 41 FDA-approved kinase inhibitors. 1; the highest positive correlation,−1; highest negative correlation. Highly correlated KIs are highlighted in red.

#### 3.1.1. VEGFR

VEGFR are essential to form and maintain blood vessel structure. VEGFR family has three members; VEGFR 1, 2 and 3. VEGFR-2 plays a major role in angiogenesis and it activates PLCγ-PKC-MAPK pathway ^43^. Renal cell carcinomas (RCC) are associated with high expression of vascular endothelial growth factor and for that reason patients with RCC are treated with VEGFR inhibitors ^44^. Among the 41 FDA-approved KIs, half of them target VEGFR with IC_50_, K_i_, K_d_ ranging from 0.01 nM to 7 μM (**Figure 7**).

**Figure 7.**
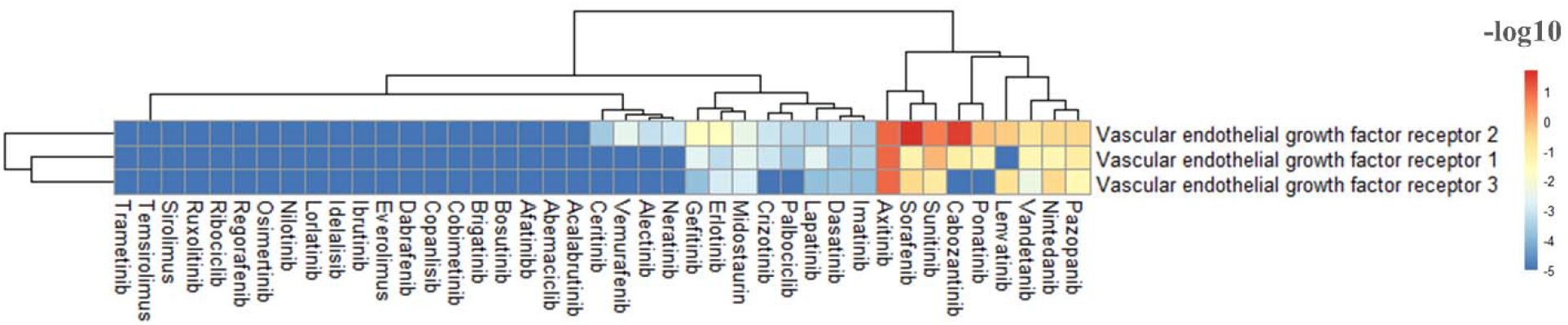
Heatmap showing which KIs target VEGFR. Value are showed in −log_10_(IC_50_, K_i_, K_d_) before applying the unbound C_max_ cutoff.

Type II-A (sorafenib, cabozantinib and axitinib) and Type II-B (sunitinib) are the strongest inhibitors of VEGFR-2 with IC_50_, K_i_ and K_d_ less than 1 nM. The difference between type II-A and B KIs is that type II-A KIs occupy the front cleft and extent to the back cleft while type II-B KIs only occupy the front cleft. Type II-A inhibitors have a residence time more than 64 min while type II-B inhibitors have less than 2.9 min ^45^. Occupying the hydrophobic back pocket enhanced the VEGFR inhibitory effect of KIs (**Figure 8**). The crystal structure of VEGFR with sorafenib, axitinib and sunitinib are available with the PDB entry of **4ASD**, **4AG8** and **4AGD** respectively. Looking at the interaction of sorafenib at the binding site, one can concludes that the amide group at the front cleft could be important to increase the activity of KIs toward VEGFR-2 and the halogen substitution at the hydrophobic back pocket could increase the sensitivity of KIs (**Figure 8**).

**Figure 8.**
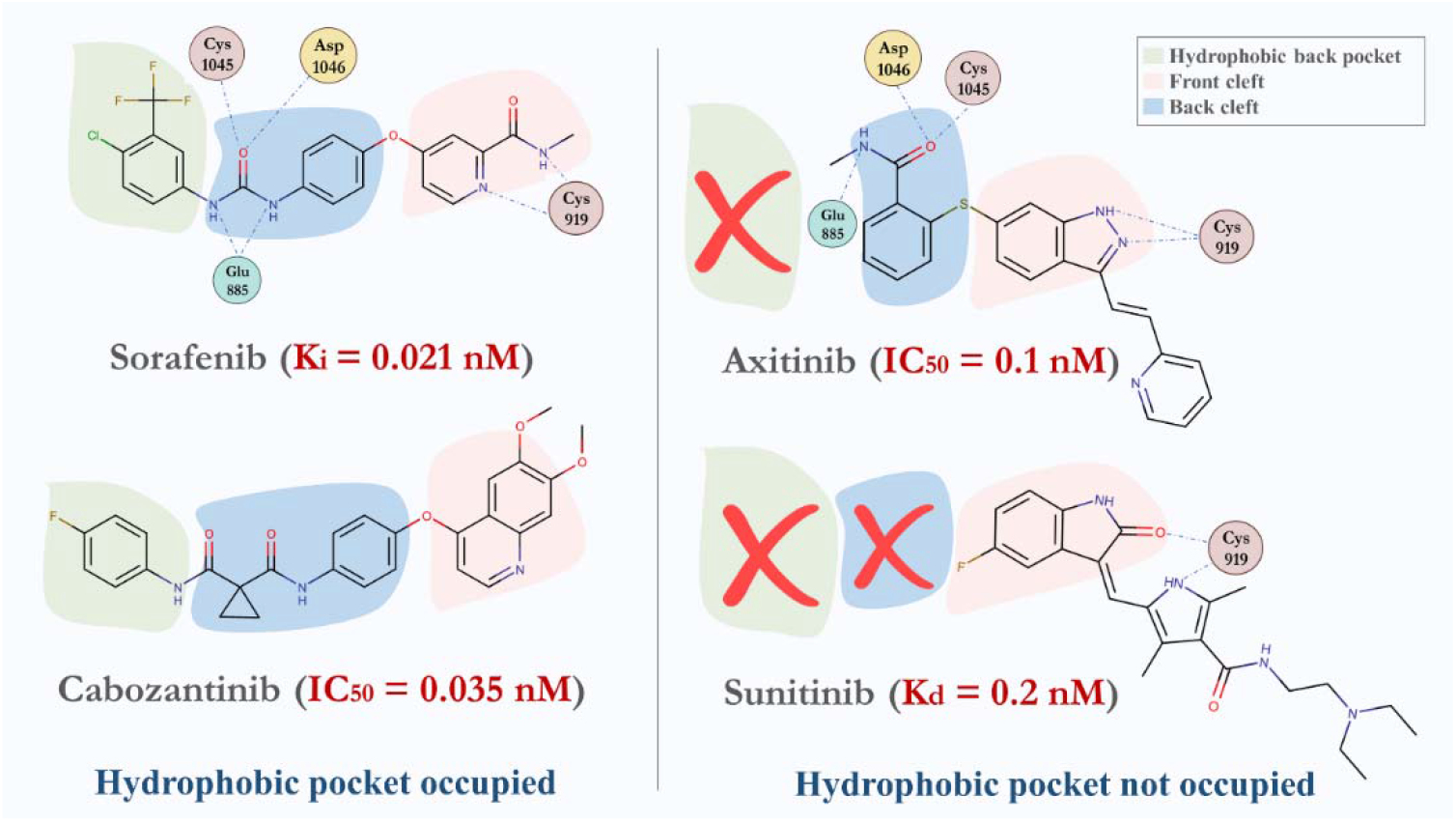
KIs of VEGFR-2 with lowest IC_50_, K_d_, K_i_. Interaction with the amino acid at the binding site of VEGFR-2 was retrieved from the PDB; sorafenib (4ASD), axitinib (4AG8) and sunitinib (4AGD). Only common interaction with the best inhibitor (sorafenib) are shown. Cabozantinib does not have a PDB entry. Blue dots represent hydrogen bonds. Sorafenib, cabozantinib and axitinib are class II-A KIs, sunitinib is class II-B KIs.

The unbound C_max_ of KIs can give an idea about the probability of binding to targets besides the primary target/s of KIs. The primary targets of sorafenib and axitinib are VEGFR 1,2 and 3. The primary targets of cabozantinib are RET, VEGFR-2 and the primary target of sunitinib is VEGFR-2. All four KIs also target stem cell growth factor receptor. By not occupying the back hydrophobic pocket; axitinib’s IC_50_ decreased 3 folds compare to cabozantinib but also increased its sensitivity toward PDGFR with K_d_ of 0.55 nM. Cabozantinib is only targeting receptor TKs, it has the highest rotatable bonds and lowest hydrogen bonds donor among the selected 4 KIs (**Figure 10C**). Other than receptor TKs, axitinib targets non-receptor kinase ABL and Aurora C. Sunitinib and sorafenib are targeting more targets compare to cabozantinib and axitinib (**Figure 9**).

**Figure 9.**
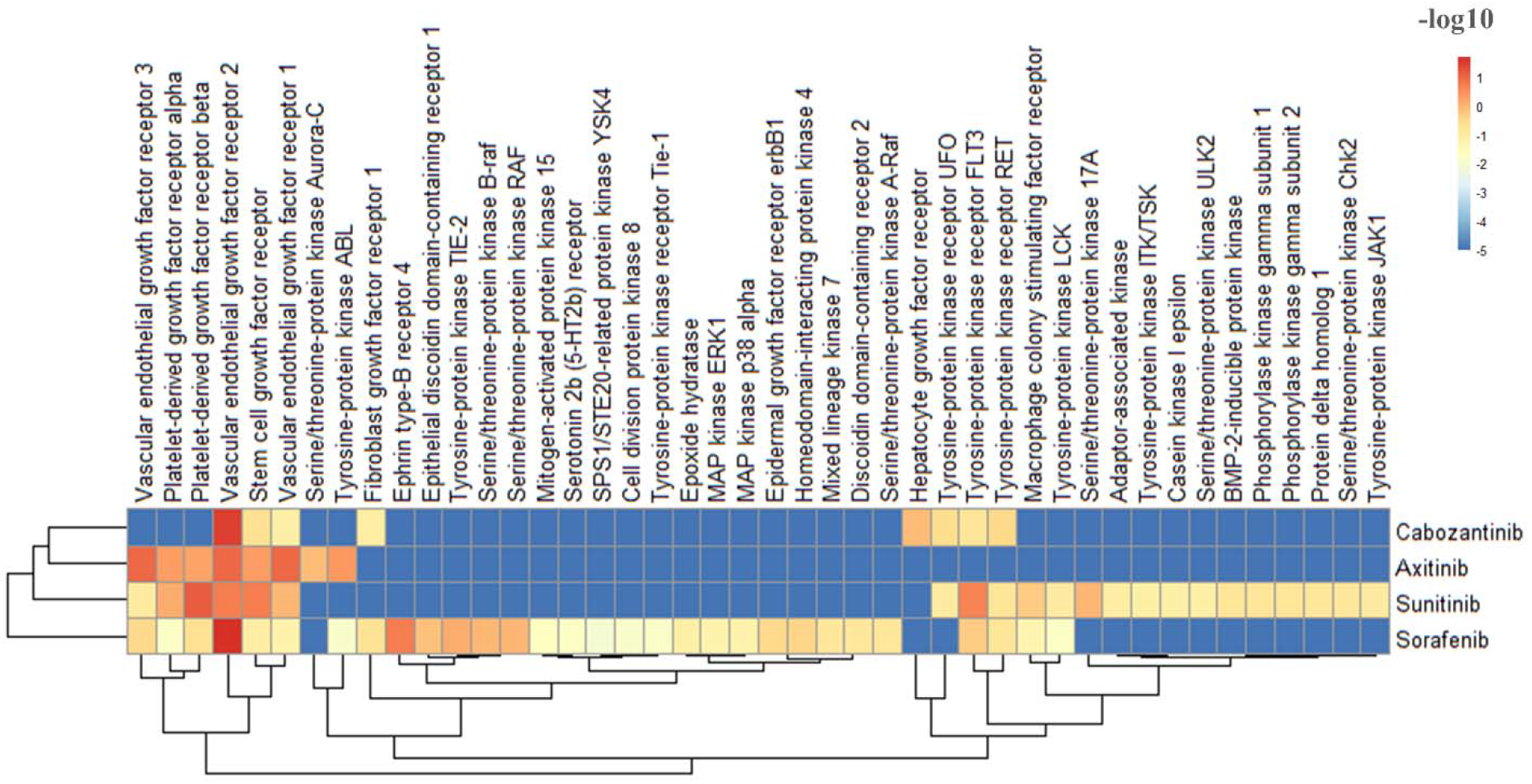
Heatmap showing the clinical targets of the VEGFR KIs (unbound C_max_). Values are showed in −log_10_(IC_50_, K_i_, K_d_).

**Figure 10.**
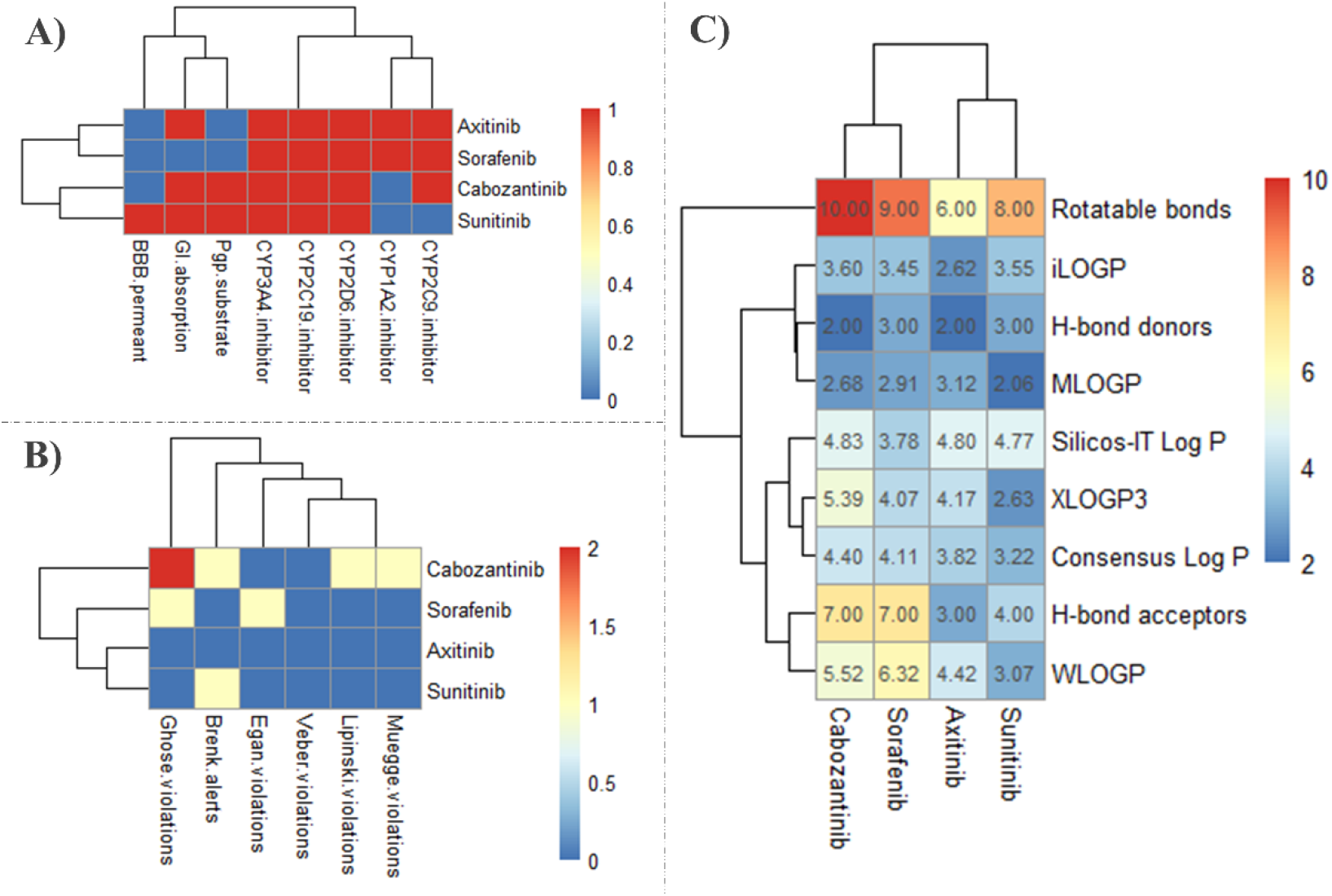
SwissADME results of VEGFR KIs. **A)** Pharmacokinetics, (yes; 1, no; 0). BBB; blood brain barrier (yes; 1, no; 0), GI; gastrointestinal (high; 1, low; 0), Pgp; P-glycoprotein, **B)** Druglikeness, **C)** Physicochemical properties & lipophilicity.

All the four VEGFR-2 KIs are FDA-approved to treat RCC. In addition, sorafenib is approved treat hepatocellular carcinoma (HCC) and differentiated thyroid cancer. Cabozantinib is approved to treat medullary thyroid cancers and HCC, sunitinib is approved to treat gastrointestinal stromal tumors (GIST) and pancreatic neuroendocrine tumors. The side effects of the KIs are related with their targets and with which cancer type/s they are used to treat. Looking at the correlation analysis in **Figure 6**, sorafenib and cabozantinib cluster next to each other, also axitinib and sunitinib cluster next to each other. Analyzing the VEGFR-2 KIs results, we noticed a positive correlation between how many targets each drug has and how many side effects have been reported for each drug (**Figure 11**). There is also a correlation between occupying the hydrophobic back pocket and causing palmar-plantar erythrodysaesthesia syndrome (PPES). The latter is a common toxic side effect of VEGFR KIs that lead to stop the treatment^46^. Out of the 41 FDA-approved KIs, 10 KIs cause PPES (**Figure S2**) and all target VEGFR with different IC_50_, K_i_ and K_d_ (**Figure 7**). According to INTSIDE (https://intside.irbbarcelona.org/index.php) ^47^, PPES is caused by interacting with DNA topoisomerase 2 or histone deacetylase 1 (HDAC1). Inhibiting HDAC decreases VEGFR-2 half-life ^48^. Sorafenib is reported to enhance the inhibition of HDAC in HCC ^49^. Other study found that inhibitors of HDAC also inhibits HGFR^50^. Out of 41 KIs, only sorafenib and cabozantinib are approved to treat HCC. In addition, cabozantinib inhibits HGFR at IC_50_ of 1.3 nM. On the other hand, VEGFR-2 KIs that not occupying the hydrophobic increase disease progression and cause serious side effect such as; neoplasm progression, renal cancer and metastatic RCC (**Figure 11**).

**Figure 11.**
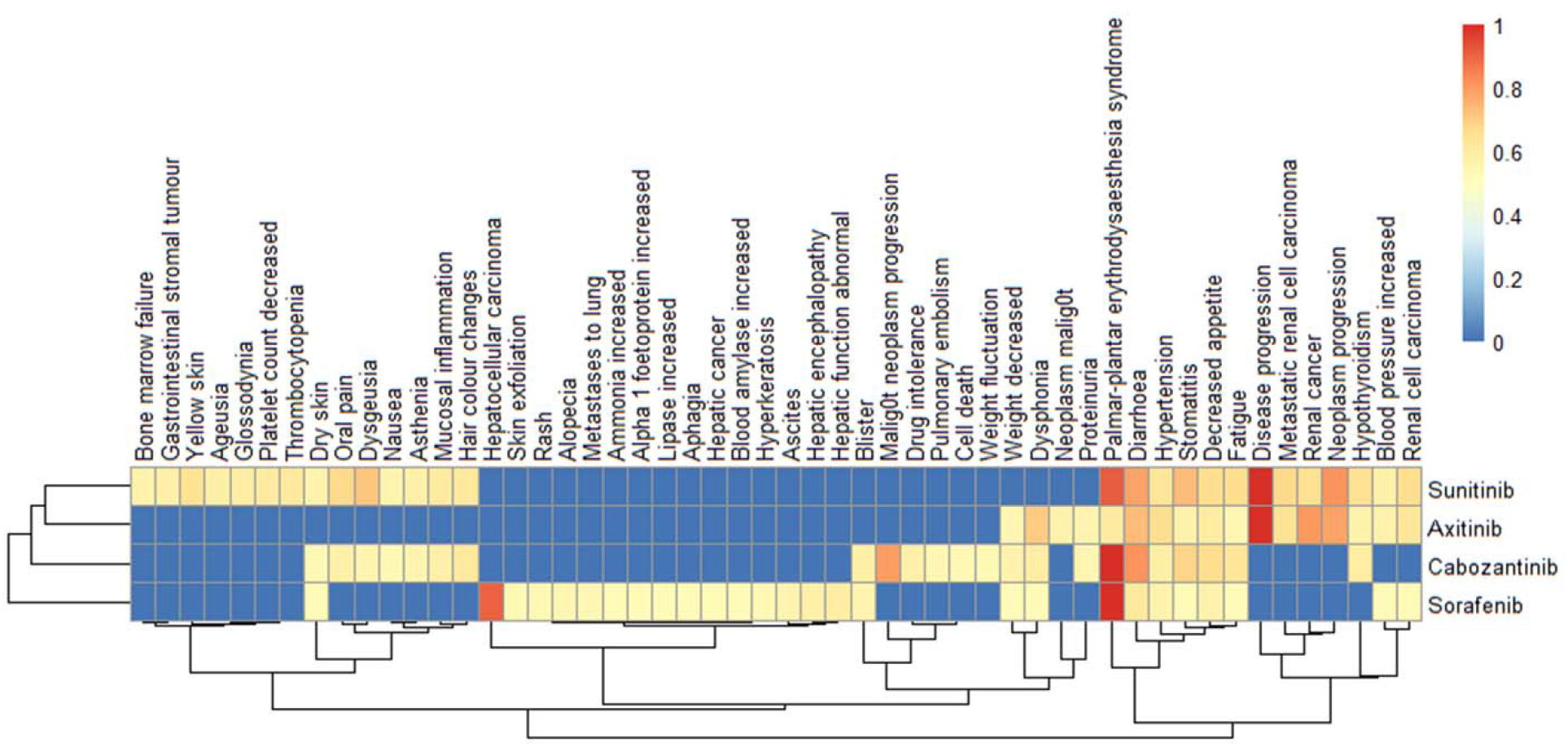
Side effect of the four KIs. **1**; highest reported side effect, **0**; side effect not reported. .

#### 3.1.2. EGFR

Dysregulation of EGFR receptors leads to the oncogenic transformation of various cell types. Upregulating EGFR has been observed in many cancer types. According to The Human Protein Atlas (https://www.proteinatlas.org/) (ATLAS), EGFR is highly expressed in glioma, colorectal, breast, prostate and lung cancer ^51^. The upregulation of EGFR is either due to L858R and T790M mutations or truncation of exon 19 at extracellular domain causing activation of the pro-oncogenic signaling pathways RAS-RAF-MEK-ERK MAPK pathway or AKT-PI3K-mTOR pathway ^52^. Afatinib, erlotinib, gefitinib are FDA-approved to treat non-small-cell lung cancer (NSCLC) while lapatinib and neratinib are FDA-approved to EGFR-(HER-2) positive breast cancers ^53^. Vandetanib is FDA-approved to treat medullary thyroid cancers ^45^. In addition, gefitinib, erlotinib, and lapatinib have been approved to treat advanced colorectal cancer, advanced NSCLC, squamous cell carcinoma of the head and neck, along with breast and pancreatic cancer ^54, 55^. Among the 41 FDA-approved drugs, half of them target EGFR with IC_50_, K_i_ and K_d_ ranging from 0.08 nM to 7 μM (**Figure 12**).

**Figure 12.**
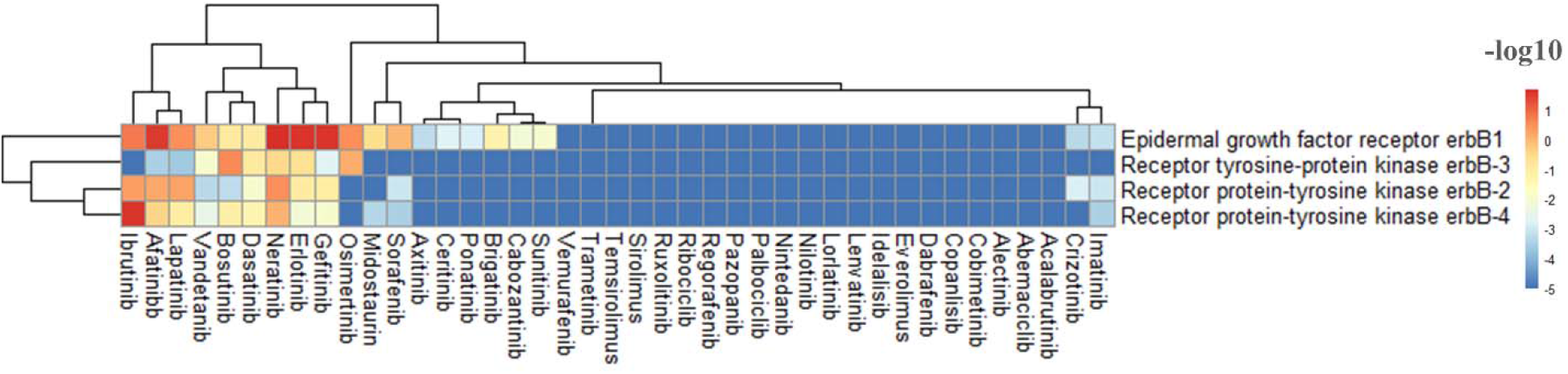
Heatmap showing which KIs target EGFR. Value are showed in −log_10_(IC_50_, K_i_, K_d_) before applying the unbound C_max_ cutoff.

Out of the selected six EGFR KIs, four (erlotinib, gefitinib, lapatinib and vandetanib) are first generation KIs and two (afatinib and neratinib) are second generation KIs. The second generation KIs can undergo Michael reaction to form a covalent bond with **–SH** of **Cys-797**. Neratinib is the strongest inhibitor of EGFR with the IC_50_ of 0.08 nM, this is due to the ability of neratinib to form a covalent bond **Cys-797** in the front pocket and it also extends deep in the hydrophobic pocket of the ATP-binding site. The K_d_ of lapatinib increased more than 5-folds compare to afatinib although it has a very similar structure to neratinib. The is because lapatinib cannot form a covalent bond with the **Cys-797**. Vandetanib’s primary targets are VEGFRs but it can inhibit EFGR with a lower K_d_ compare to VEGFR (**Figure 13**).

**Figure 13.**
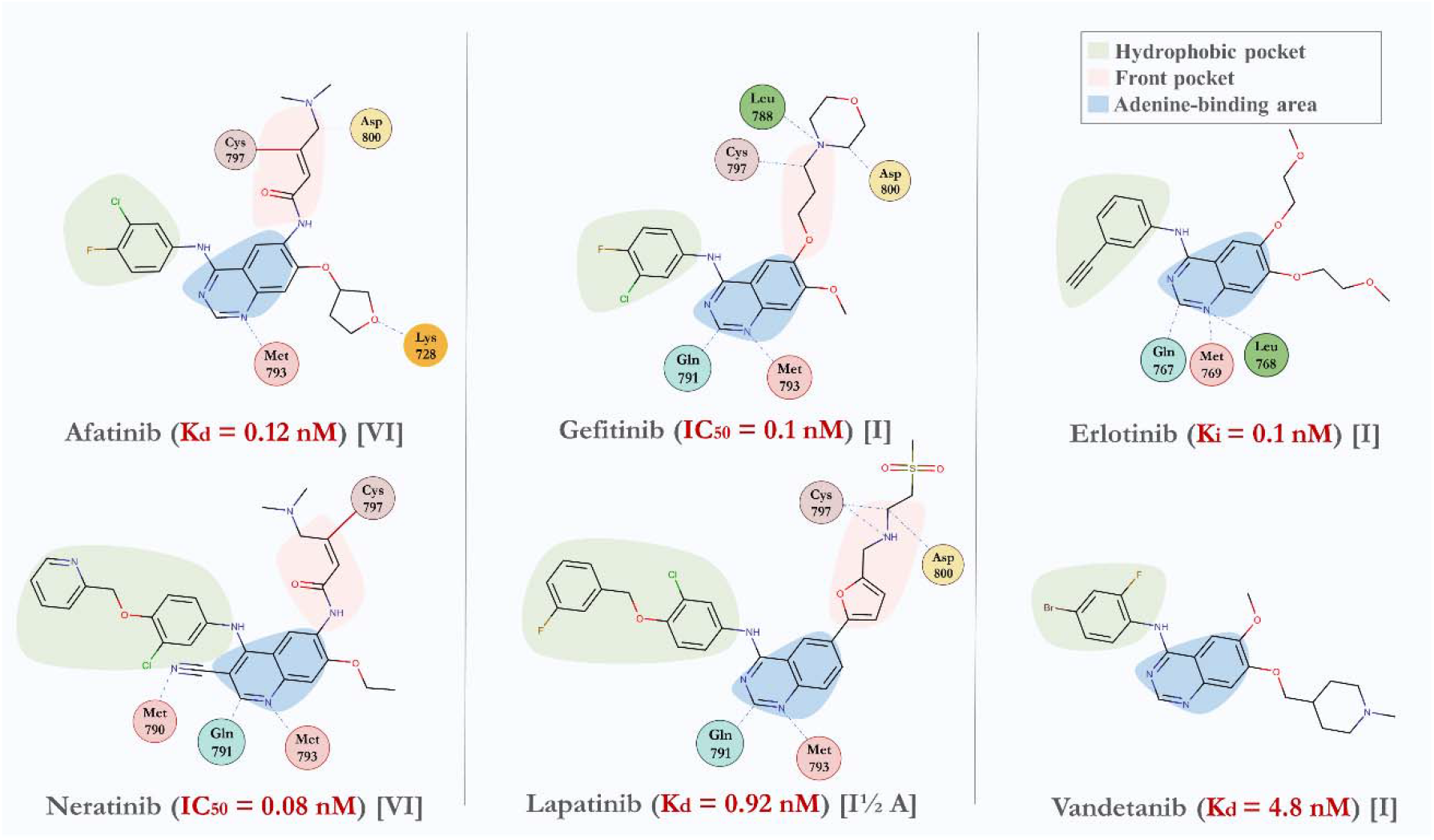
KIs of EGFR with lowest IC_50_, K_d_, K_i_. Interaction with the amino acid at the binding site of EGFR was retrieved from the PDB; afatinib (**4G5J**), neratinib (**2JIV**), gefitinib (**4I22**), lapatinib (**1XKK**), erlotinib (**4HJO**). Vandetanib does not have a PDB entry for EGFR. Interaction at the hydrophobic pocket are not shown. Blue dots represent hydrogen bonds, dark-red lines represent covalent bonds.

We calculated the unbound C_max_ of the selected six EGFR KIs and only selected the targets that are within the range of unbound C_max_. Lapatinib and afatinib are only targeting the EGFR family. Neratinib is able also to target the mitogen-activated protein kinase kinase kinase kinase 5 (MKKK5) with K_d_ of 0.65 nM. Vandetanib has the most targets followed by erlotinib and gefitinib (**Figure 14**). Vandetanib has the lowest rotatable bonds 6 (highest K_d_) compare to the other EGFR KIs and neratinib has the highest rotatable bonds 12 (lowest IC_50_) (**Figure 15**). This could contribute, along with other factors, to the sensitivity/selectivity of KIs to EGFR.

**Figure 14.**
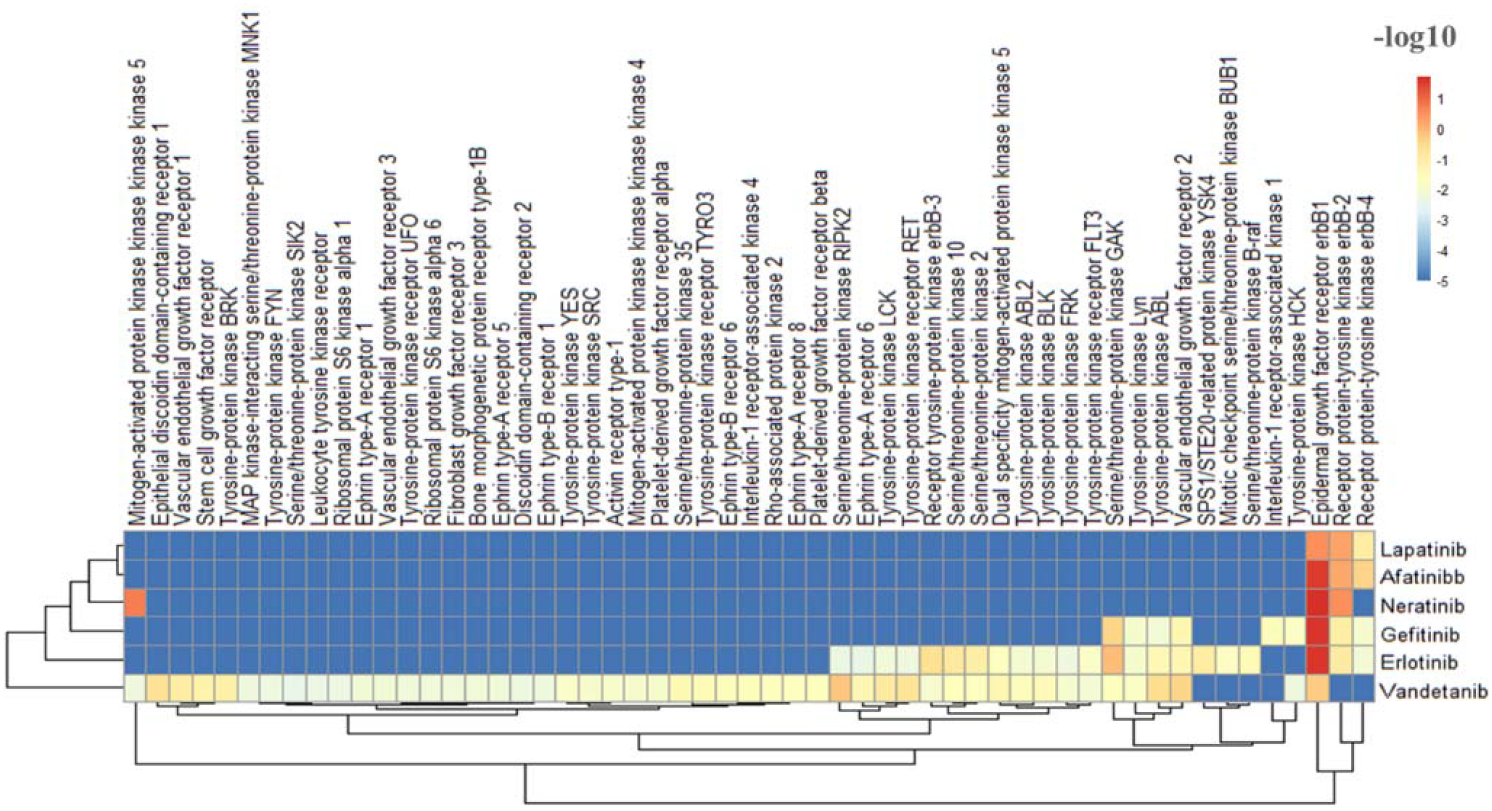
Heatmap showing the clinical targets of the EFGR KIs (unbound C_max_). Values are showed in −log_10_ (IC_50_, K_i_, K_d_).

**Figure 15.**
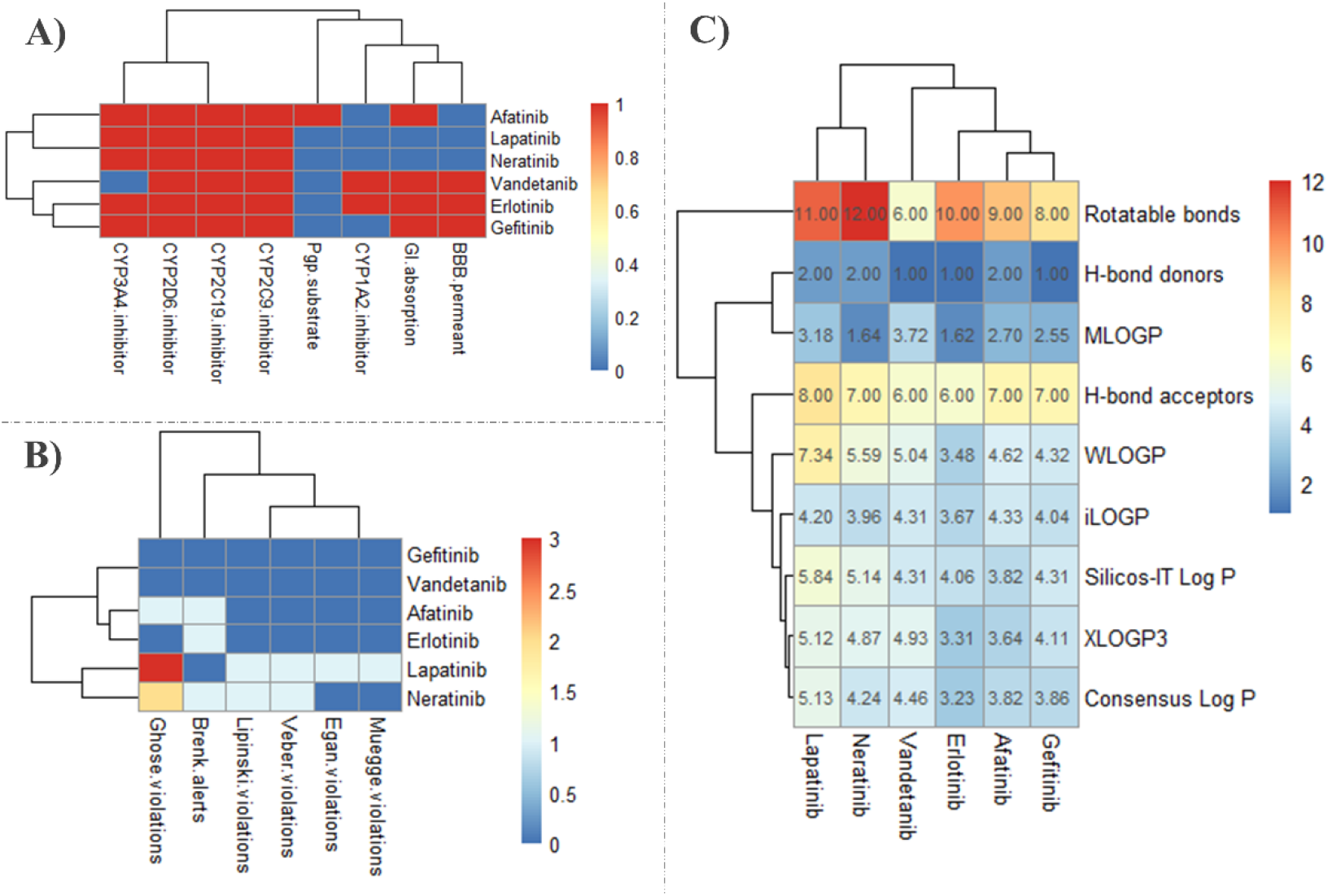
SwissADME results of EGFR KIs. **A)** Pharmacokinetics, (yes; 1, no; 0). BBB; blood brain barrier (yes; 1, no; 0), GI; gastrointestinal (high; 1, low; 0), Pgp; P-glycoprotein, **B)** Druglikeness, **C)** Physicochemical properties & lipophilicity.

Looking at the side effects, forming a covalent bond or/and extending further into the hydrophobic pocket increase the side effect of the KIs. Since vandetanib is used to treat medullary thyroid cancers, the reported side effects are different than the others EGFR KIs. Erlotinib most reported side effect is rash (**Figure 16**) and in fact it is the most rash-causing KI among the 41 KIs studied here (**Figure S2**). It has been reported that rash is induced by EGFR Inhibitors^56^ but not all EGFR KIs cause rashes (**Figure 16**). It appears that other target/s may contribute to this side effect. Analyzing the 41 KIs target and their side effects, it looks like serine/threonine-protein kinase B-raf is the common target among the KIs that cause rashes. Rash is mostly reported in patients under treatment with serine/threonine-protein kinase B-raf inhibitors (>70% with vemurafenib)^57^. Vemurafenib is the second KIs that causes rash after erlotinib (**Figure S2**). Among the EGFR KIs, only erlotinib inhibits B-raf, it is therefore intriguing to study whether the increased rash observed with erlotinib treatment could be due to its dual inhibition of EGRF and B-raf at clinically relevant doses. Another serious side effect caused by EGFR KIs is malignant neoplasm progression. Target validation database reported EGFR as the first target (with the highest score) out of 5858 targets associated with high grade malignant neoplasm ^42^. Malignant neoplasm progression was reported the highest in patients treated with afatinib and gefitinib but interestingly it was not reported in patients using neratinib and vandetanib. In one clinical trial, both afatinib and gefitinib caused death due to malignant neoplasm progression^58^. Looking at the targets of EGFR KIs, neratinib and vandetanib are the only EGFR KIs that do not target EGFR-4 (**Figure 14**). This therefore places EGFR-4 into a candidate position relating to malignant neoplasm progression. The only similarity between afatinib and gefitinib is that both have two halogens substitution (Cl and F) next to each other in the aromatic ring that interact with the hydrophobic pocket (**Figure 16**). Additionally, all patients treated with EGFR KIs suffer from diarrhea. According to INTSIDE; erlotinib, gefitinib and lapatinib cause diarrhea by affecting CYP3A4 and multidrug resistance protein. SwissADME predicted all EGFR KIs except vandetanib (lowest reports of diarrhea) could inhibit CYP3A4 (Figure 15A).

**Figure 16.**
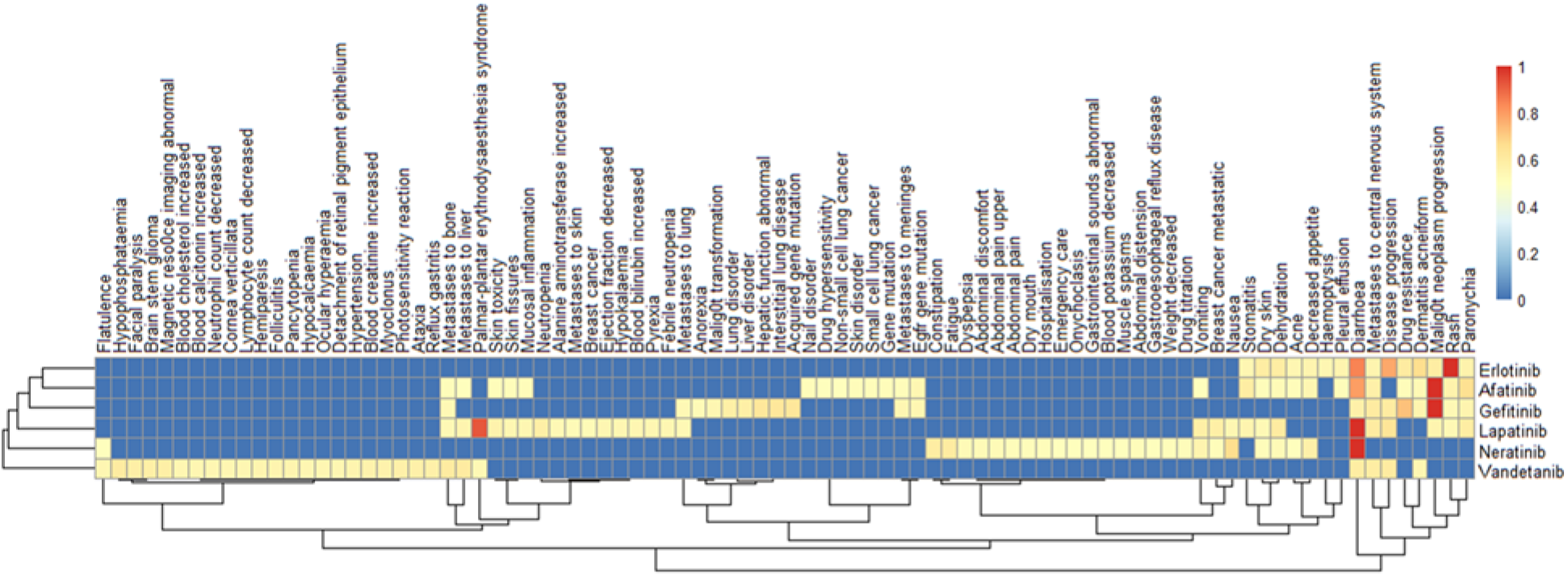
Side effect of the EFGR KIs. **1**; highest reported side effect, **0**; side effect not reported.

#### 3.1.3. ALK receptor

According to the World Health Organization (WHO), lung cancer is the leading cause of cancer-related death, in 2018 lung cancer caused more than 1.70 million deaths worldwide. 85% of lung cancer are NSCLC ^59^, approximately 5%^60^ of NSCLC are ALK positive. ALK receptor is a part of a signaling pathways that control proliferation, migration and differentiation^61^. ALK positive cancer can be solid and non-solid cancer. ALK positive NSCLC is caused when ALK gene in the DNA is combined/fused with a neighbor gene called EML4 forming a new carcinogenic version of ALK protein; EML4-ALK. Another ALK fusion protein is NPM-ALK and it occurs in anaplastic large cell lymphoma (ALCL) ^62^. ALK receptor like any other TK receptor is located at the cell surface and its activated by a ligand, but a fusion protein such as EML4-ALK remains intracellular and its constantly active (ALK is activated by the dimerization with its partner, EML4) and this triggers several signaling pathway such as STAT3, ERK, PLCγ, and PI3K/Akt pathways ^62^. Another similar mutation in NSCLC is ROS1 rearrangement. It occurs in 2% of NSCLC. ROS1 is a receptor TK that is related to ALK. ROS1-positive NSCLC is caused due to the fusion of ROS1 gene with a neighbor gene and it’s one of the aggressive cancers that can metastasis to the brain and the bones ^63^. According to ATLAS, ALK receptor is also expressed in colorectal, breast, prostate, and skin cancer. Ceritinib, crizotinib and lorlatinib are very strong inhibitors of ALK (**Figure 18**) and all of them are FDA-approved to treat ALK-positive NSCLC. It is noted that lorlatinib is the strongest inhibitor of proto-oncogene TK ROS (**Figure 17**) with a K_i_ of 0.1 nM compare to ceritinib (0.4 nM), crizotinib (0.6 nM) but only crizotinib is approved to treat ROS1-postive NSCLC. Looking at the whole targets of the three ALK KIs (unbound C_max_), crizotinib tends to have more targets than ceritinib and lorlatinib and it inhibits HGFR at Ki of 0.2 nM (**Figure 19**). In fact, it’s one of the two KIs that inhibit HGFR along with cabozantinib (IC_50_ = 1.3 nM) (**Figure 3**). Additionally, crizotinib targets the homologs of yeast Sterile 7, 11, 20 kinases (STE kinases) (**Figure 19**).

**Figure 17.**
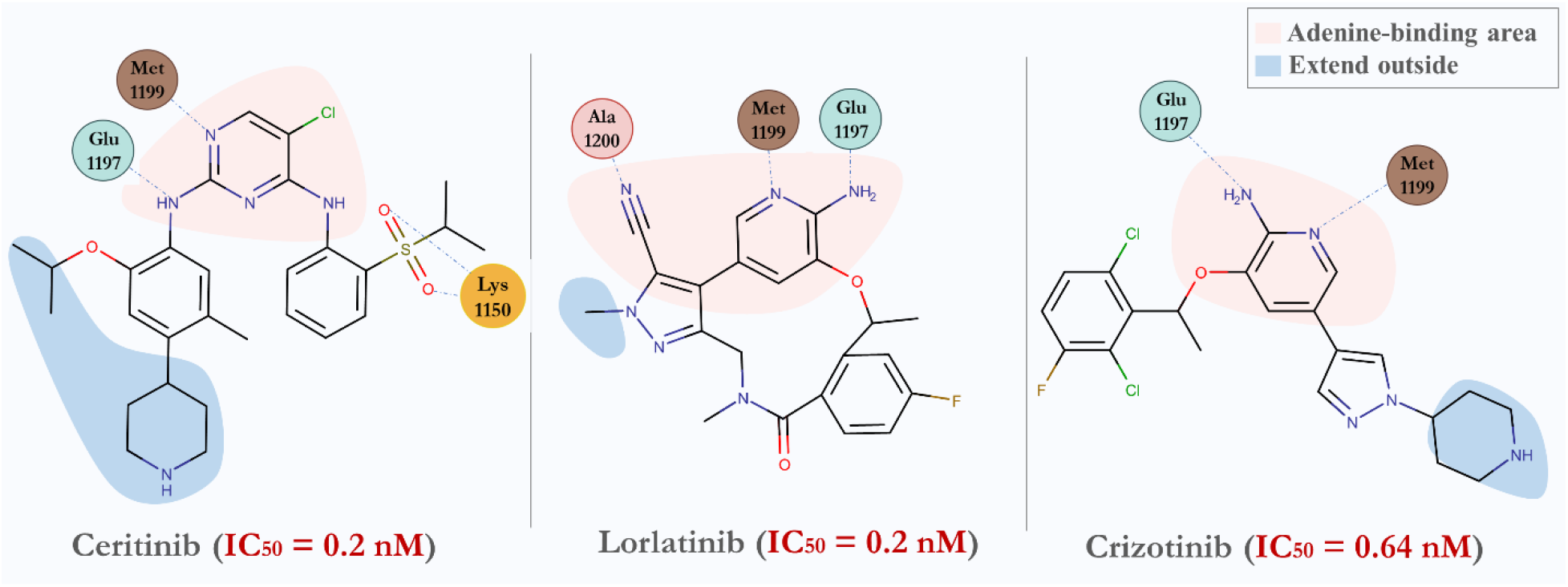
KIs of ALK with lowest IC_50_. Interaction with the amino acid at the binding site of ALK was retrieved from the PDB. Ceritinib **(4MKC)**, lorlatinib **(5AA8)** and crizotinib **(2XP2)**. Blue dots represent hydrogen bonds.

**Figure 18.**
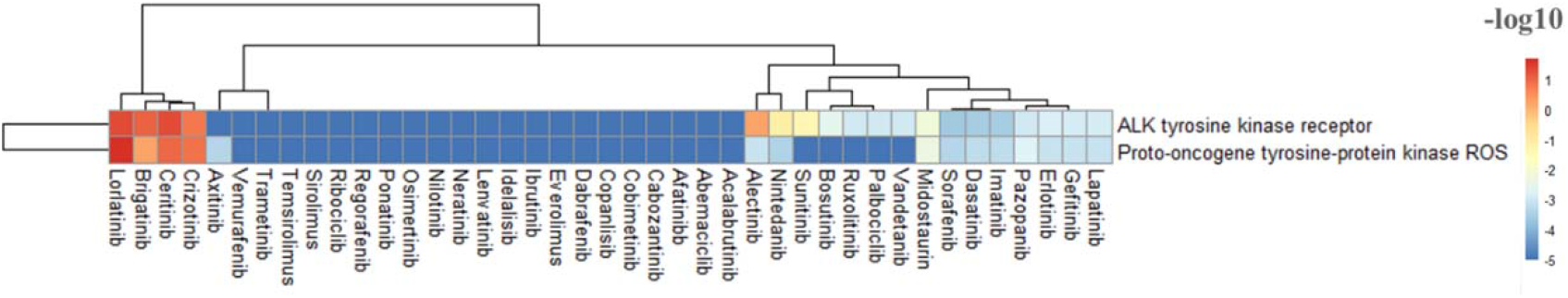
Heatmap showing KIs that target ALK and ROS receptor TKs. Value are shown in −log_10_(IC_50_, K_i_, K_d_) before applying the unbound C_max_ cutoff.

**Figure 19.**
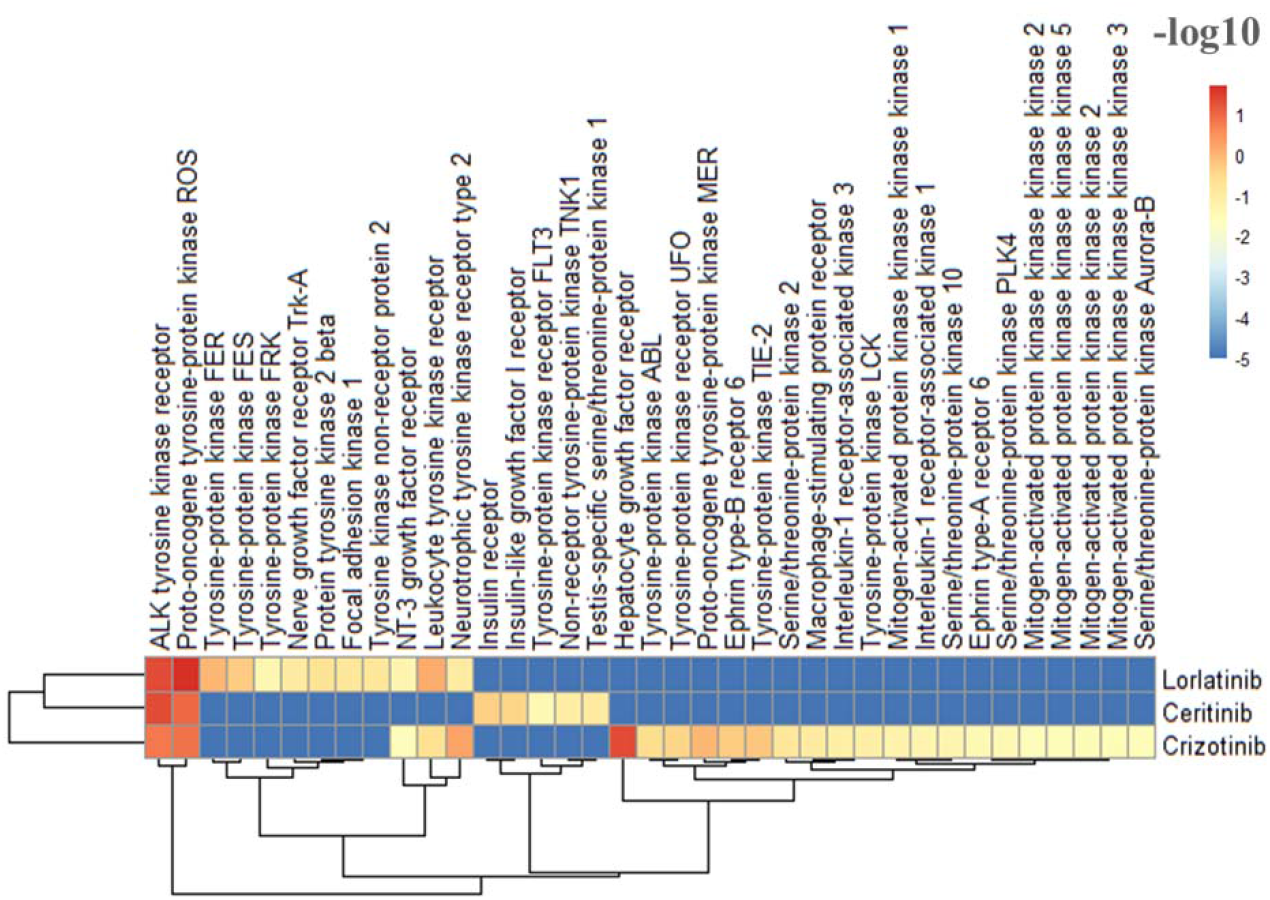
Heatmap showing the clinical targets of the ALK KIs (unbound C_max_). Values are shown in −log_10_ (IC_50_, K_i_, K_d_).

Ceritinib has the highest side effects and most of those side effects are not reported in lorlatinib and crizotinib, this could be related to high number (9) rotatable bonds in ceritinib which could impart higher conformational flexibility in ceritinib structure (**Figure 20**). All the three ALK KIs cause neoplasm progression with ceritinib causing also malignant neoplasm progression (highest reported) (**Figure 21**). Ceritinib cause some heart problems as side effect e.g.; inflammation of the pericardium (pericarditis), electrocardiogram QT prolonged and pericardial effusion. An in silico study identified the non-receptor TNK1 to be a myocardial infarction-related protein ^64^, ceritinib is the only ALK KIs that inhibits TNK1 (K_d_=27 nM). In addition, ceritinib causes abnormal increase in some hepatic enzymes (alanine aminotransferase (> 5 times), gamma-glutamyltransferase and aspartate aminotransferase) which leads to severe hepatotoxicity ^65^. Both ceritinib and lorlatinib increase patients’ lipid profile.

**Figure 20.**
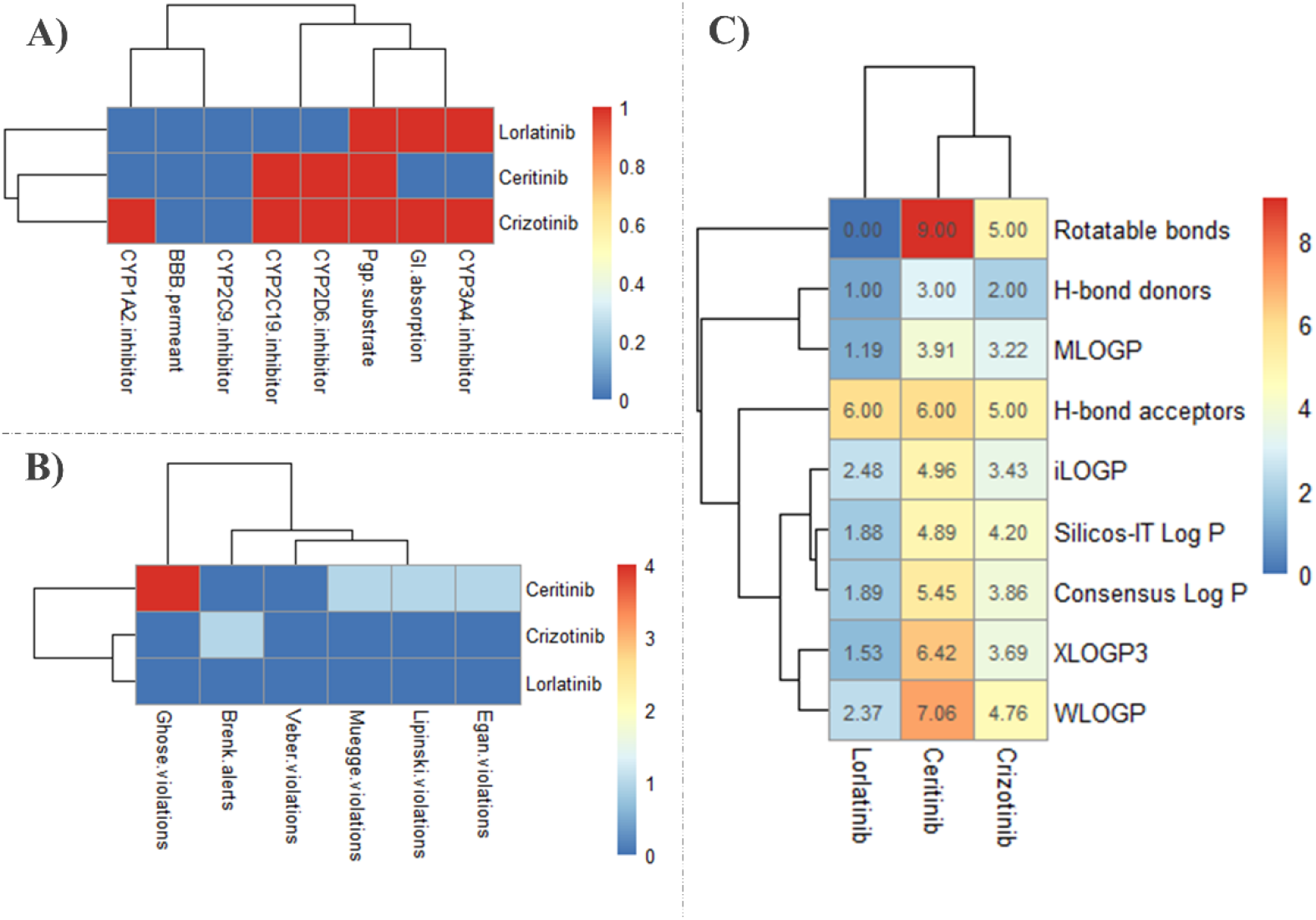
SwissADME results of ALK KIs. **A)** Pharmacokinetics, (yes; 1, no; 0). BBB; blood brain barrier (yes; 1, no; 0), GI; gastrointestinal (high; 1, low; 0), Pgp; P-glycoprotein, **B)** Druglikeness, **C)** Physicochemical properties & lipophilicity.

**Figure 21.**
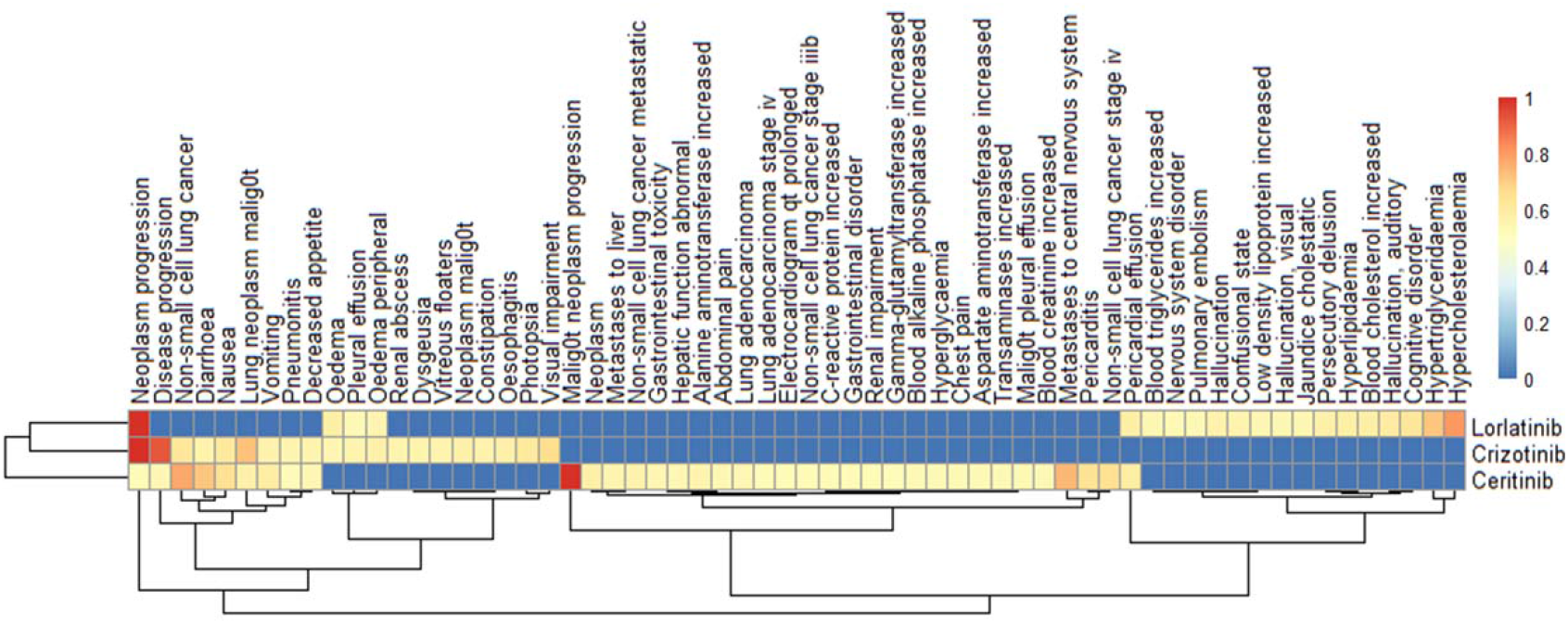
Side effect of the ALK KIs. 1; highest reported side effect, **0**; side effect not reported.

#### 3.1.4. ABL family

ABL is a non-receptor tyrosine-protein kinase that plays an important role in cell proliferation, migration and survival ^3^. It has two forms; ABL1 and ABL2. ABL1 has a DNA binding domain (for DNA damage-repair) while ABL2 has the ability to bind to microtubules and actin (for cytoskeletal remodeling functions) ^66^. ABL protein kinase is well-known because of the BCR-ABL mutation, the mutation happens when part of breakpoint cluster region protein (BCR) gene breakdown from chromosome 9 (BCR original location) and moved to ABL genes lactation on chromosome 22 ^67^. The new formed gene is called philadelphia chromosome and this gene is responsible for the production BCR-ABL fusion protein. This type of mutation elevates tyrosine kinase activity and cause cells to divide uncontrollably, and it exists in certain types of leukemia such as chronic myeloid leukemia (CML) and acute lymphoblastic leukemia (CLL) ^68^. In 70 % of patients the blasts are myeloid, and in 30% they are lymphoid ^69^. KIs such as bosutinib, dasatinib, imatinib, nilotinib and ponatinib inhibit proliferation and induces apoptosis in Bcr-Abl positive cell lines and they are used to treat many types of cancer including leukemia. Among the 41 FDA-approved KIs, more than half of them target ABL with IC_50_ ranging from 0.019 nM (dasatinib) to 6 μM (neratinib) (**Figure 22**).

**Figure 22.**
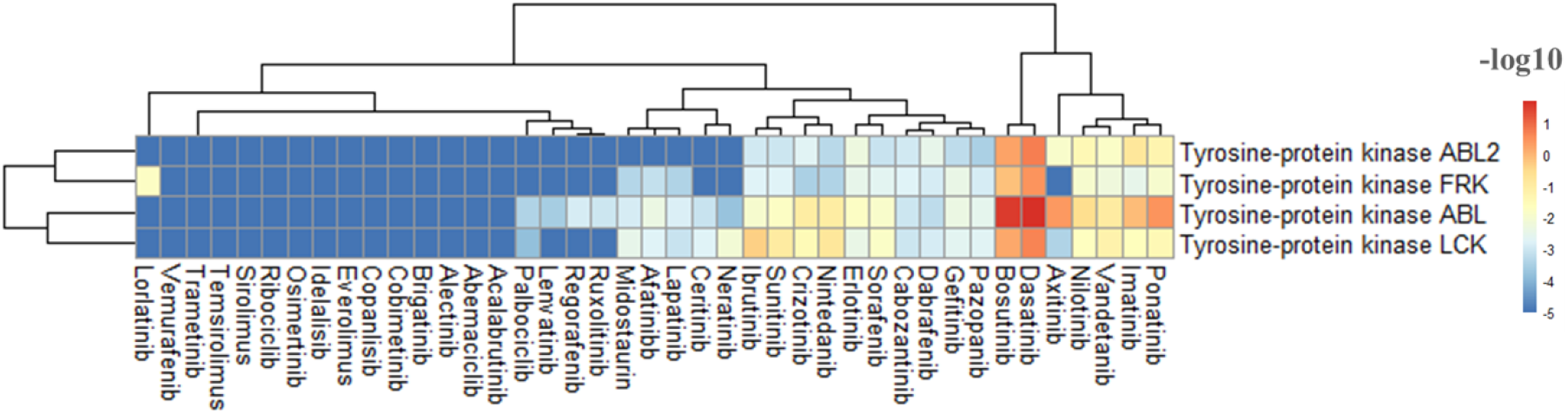
Heatmap showing which KIs target ABL family, LCK and FRK. Value are showed in −log_10_(IC_50_, K_i_, K_d_) before applying the unbound C_max_ cutoff.

Dasatinib and bosutinib are FDA-approved to treat CML. Ponatinib and nilotinib are approved to treat Ph+ CML. On the other hand, imatinib is approved to treat several cancer types (CLL, Ph+ CML, ALL mantle cell lymphomas, chronic eosinophilic leukemias). Only dasatinib binds to the active ABL (type I KIs), while, ponatinib, imatinib and nilotinib bind to the inactive ABL and extend to the back cleft (type IIA KIs). Bosutinib binds to the inactive ABL but do not extend to th back cleft (type IIB KIs) (**Figure 23**).

**Figure 23.**
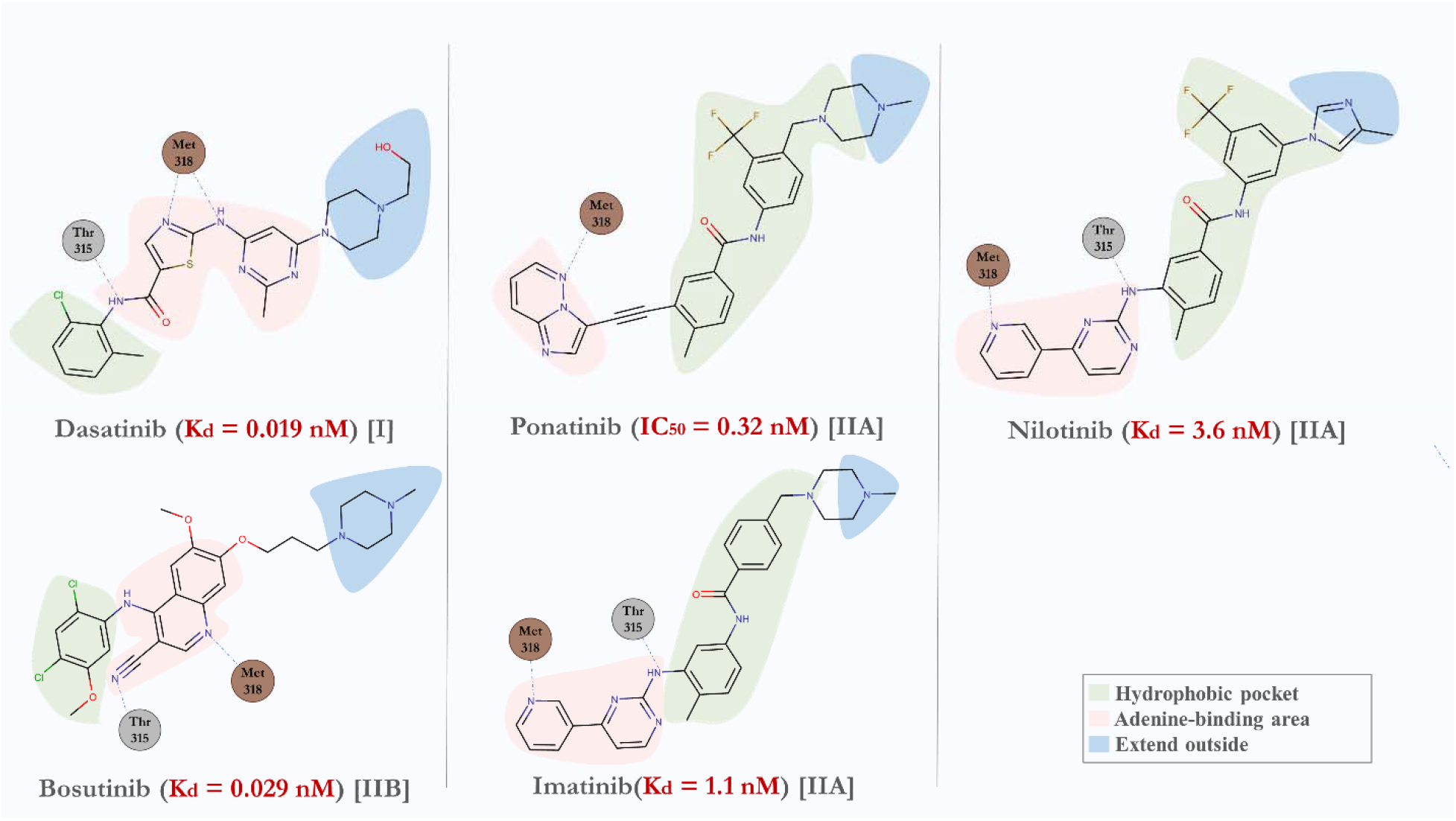
KIs of ABL with lowest IC_50_, K_d_. Interaction with the amino acid at the binding site of ABL was retrieved from the PDB. Dasatinib **(2GQG)**, bosutinib **(3UE4)**, ponatinib **(3OXZ)**, imatinib **(1IEP)** and nilotinib **(3CS9)**. Blue dots represent hydrogen bonds.

All ABL KIs also inhibit TK LCK (**Figure 24**). Dasatinib and bosutinib inhibit > 25 similar targets. On the other hand, imatinib and nilotinib share similar part of their structure (4-(Pyridin-3-yl)pyrimidin-2-amine). This structural similarity correlates with some targets only imatinib and nilotinib inhibit. Among the ABL KIs, only imatinib and nilotinib can inhibit carbonic anhydrase family with K_i_ ranging from 4.1 to 468 nM and only imatinib can target serotonin receptor 2A (Ki=540 nM) and serotonin transporter (K_i_= 745 nM) (**Figure 24**).

**Figure 24.**
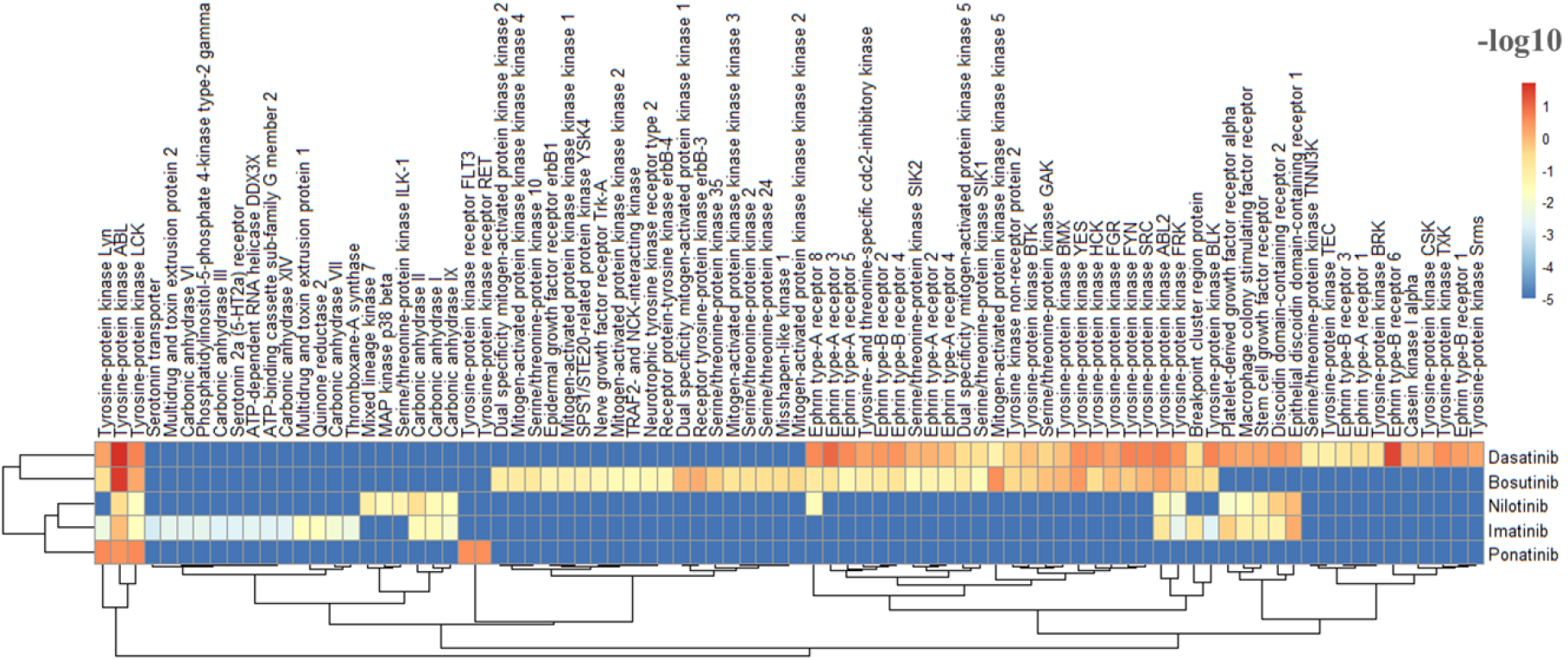
Heatmap showing the clinical targets of the ABL KIs (unbound C_max_). Values are showed in −log10 (IC_50_, K_i_, K_d_).

**Figure 25.**
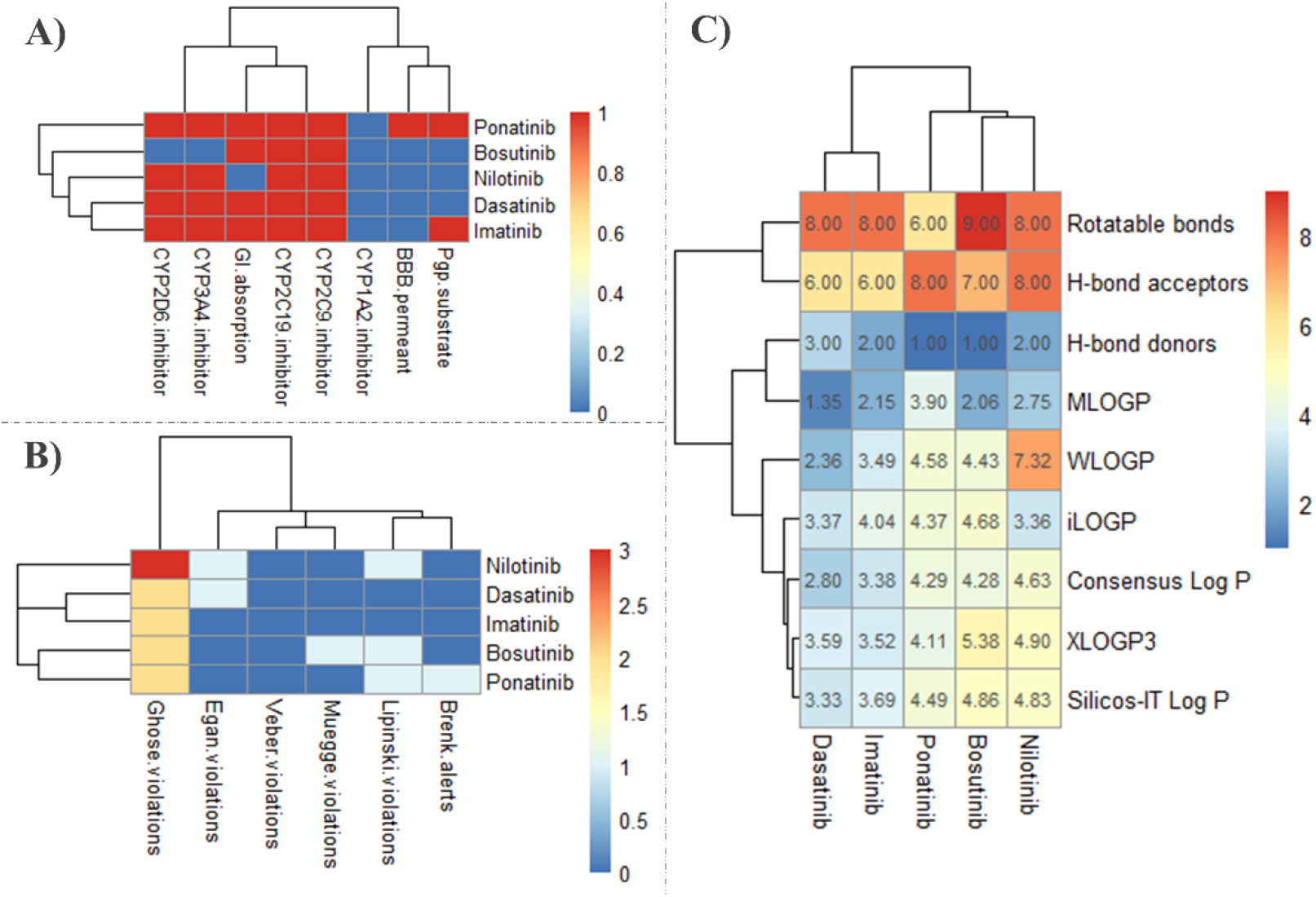
SwissADME results of ABL KIs. **A)** Pharmacokinetics, (yes; 1, no; 0). BBB; blood brain barrier (yes; 1, no; 0), GI; gastrointestinal (high; 1, low; 0), Pgp; P-glycoprotein, **B)** Druglikeness, **C)** Physicochemical properties & lipophilicity.

All ABL KIs cause cytogenetic (functioning of chromosomes) abnormalities and pleural effusion (**Figure 26**). According to INTSIDE, pleural effusion is caused by ABL KIs by targeting two protein kinases; ABL1 and stem cell growth factor KIT (**Figure 27**). Dasatinib has the highest pleural effusion reports^70^ and this probably due to its strong inhibitory effect on both ABL and stem cell growth factor receptor (K_d_= 0.57 nM). Peripheral arterial occlusive disease happens only in CML patients treated with nilotinib^71^ and ponatinib ^72^. This is probably due to the three fluorine substations in the aromatic ring.

**Figure 26.**
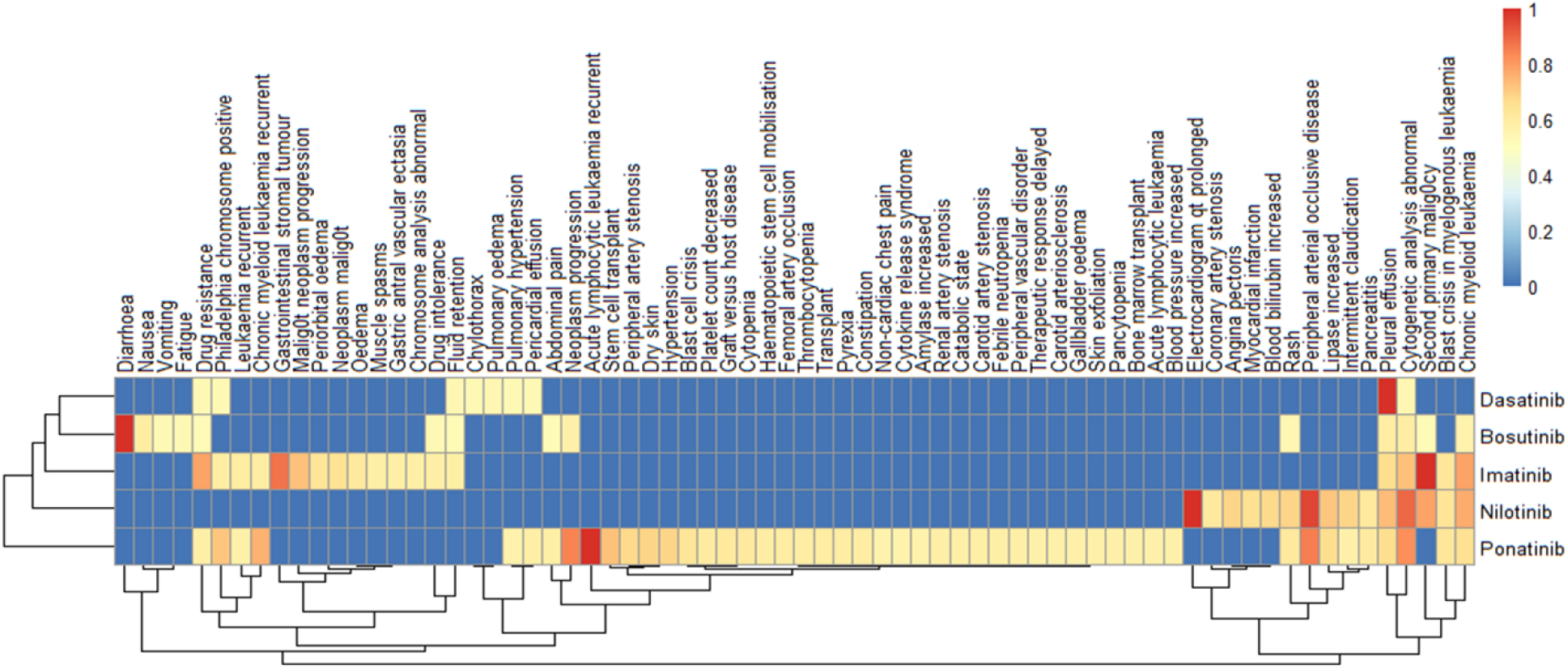
Side effect of the ABL KIs. _1_; highest reported side effect, _0_; side effect not reported

**Figure 27.**
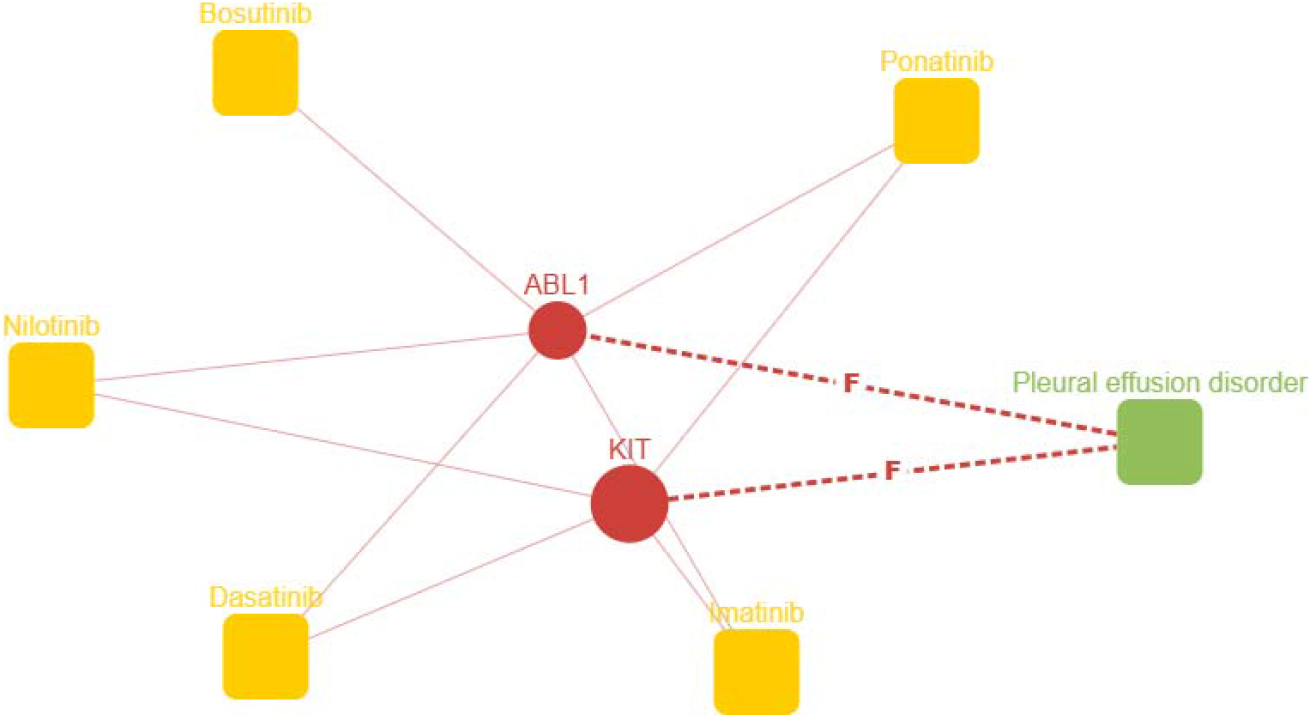
Target network related to pleural effusion disorder. F; Over-represented protein Bonferroni p-value: 1.0e-5.

## 4. CONCLUSION

We have analyzed the molecular targets of 41 U.S. FDA-approved KIs. We have chosen 18 drugs (Tyrosine KIs) for further analysis to identify their possibly engagement in vivo in clinically relevant doses and whether there can be any correlation between their targets and the reported side effects. To explore all clinical targets of the 18 KIs, we retrieved data from ChemBL and other public databases to obtain or calculate the free (unbound) Cmax of each KI. Considering known efficacy or affinity metrics (IC_50_, K_i_ and K_d_), we only chose targets for which the cognate affinities lie within the reported free Cmax values, thereby allowing plausible interaction in clinical doses. Our analyses seem to have revealed some interesting correlation. For example, we found a correlation between occupying the hydrophobic back pocket of VEGFR (sorafinib, cabozantinib) and causing PPES by inhibiting HDAC. It appears that targeting EGFR and serine/threonine-protein kinase B-raf increase rash in cancer patients treated with erlotinib, in fact erlotinib is the highest KIs with reported rash among the 41 KIs. We also concluded that EGFR-4 maybe a candidate that cause malignant neoplasm progression. Also, the two halogens substitution (Cl and F) next to each other in the aromatic ring that interact with the hydrophobic pocket of EGFR (afatinib, gefitinib) could be the reasons why afatinib and gefitinib are the highest KIs with malignant neoplasm progression reports. Ceritinib has the highest side effects and most of those side effects are not reported in lorlatinib and crizotinib, this could be related to high number (9) rotatable bonds in ceritinib which could impart higher conformational flexibility in ceritinib structure. Ceritinib also cause some heart problems as side effect e.g.; inflammation of the pericardium (pericarditis), electrocardiogram QT prolonged and this could be by inhibiting the non-receptor TNK1. The ABL KIs, imatinib and nilotinib share similar part of their structure (4-(Pyridin-3-yl)pyrimidin-2-amine). This structural similarity correlates with some targets only imatinib and nilotinib could interact and inhibit (for example, the carbonic anhydrase family). These correlations we found may in future help to formulate interesting hypothesis for explaining or predicting relative efficacy, selectivity and safety of these approved inhibitors and prompt appropriate experimental approaches towards validation of such correlations.

## ASSOCIATED CONTENT

### Supporting Information

Molecular targets of 12 kinase inhibitors, Skin related side effect of the kinase inhibitors, Kinase inhibitors that cause metastases and maliganant neoplasm, FDA-approved drugs that pass blood brain barrier, Complete list of the side effects of 41 kinase inhibitors.

## AUTHOR INFORMATION

### Notes

The authors declare no competing financial interest.

## ABBREVIATIONS USED

ATLAS: The Human Protein Atlas
C_max_: maximum serum concentration
FDA: U.S. Food and Drug Administration
HDAC1: histone deacetylase 1
IC_50_: half-maximal inhibitory concentration
K_i_: inhibition constant
K_d_: mean dissociation constant
KIs: Kinase Inhibitors
LL: Log likehood
MKKK5: mitogenactivated protein kinase kinase kinase kinase 5
PPES: palmar-plantar erythrodysaesthesia syndrome
TK: Tyrosine Kinase

